# Dysregulation of innate immunity and cellular metabolism through virus-induced deISGylation

**DOI:** 10.1101/2025.09.08.674928

**Authors:** Junji Zhu, GuanQun Liu, Jielin Xu, Kun Li, Christopher M. Goins, Huaxu Yu, Zuberwasim Sayyad, Yadi Zhou, Evangeline White, Oliver Fiehn, Shaun R. Stauffer, Feixiong Cheng, Michaela U. Gack

## Abstract

Interferon-stimulated gene 15 (ISG15) regulates diverse cellular responses including antiviral immunity through its conjugation to proteins, a process known as ISGylation. Several pathogens, including SARS-CoV-2, subvert ISGylation by encoding deISGylating enzymes. However, the direct targets and physiological consequences of coronaviral deISGylation remain poorly defined. Here, we ablated the deISGylating activity of the SARS-CoV-2 papain-like protease (PLpro) and found that loss of deISGylation boosted innate immune activation, attenuated virus replication, and promoted viral clearance in human cells and in mice. Through untargeted metabolomics and ISGylome proteomics analyses, combined with functional studies, we discovered in molecular detail how the activities of key metabolic enzymes in glycolysis, the pentose phosphate pathway, and oxidative stress are controlled by PLpro deISGylation. These findings provide fundamental new insight into how reversible ISGylation regulates immunity and metabolic processes at the molecular level and highlight viral deISGylation as a major viral tactic for rewiring immunometabolism.

## INTRODUCTION

ISGylation is a reversible process in which ISG15, a ubiquitin-like (UBL) protein, is covalently attached to target proteins to regulate their interactome, stability, subcellular localization and/or signaling ability^1,2^. As ISG15 and the components of its conjugation machinery (i.e., E1, E2 and E3 enzymes) are inducible by interferons (IFNs), ISGylation is sharply increased during pathogen infection or in non-infectious inflammatory conditions where cytokine production is uncontrolled. Infection-triggered ISGylation of viral proteins is part of the cell-intrinsic immune response and typically leads to inhibition of viral protein function and suppression of virus replication^3,4^. On the other hand, recent research showed that HERC5-mediated ISGylation promotes innate immune defenses by activating immune sensor or signaling proteins such as MDA5, cGAS, STING and IRF3^5–8^. In turn, many viral pathogens employ tactics to reverse ISGylation events in infected cells to restore viral protein function and/or suppress innate immunity, overcoming virus restriction^3,4^.

Coronaviruses such as severe acute respiratory syndrome coronavirus 2 (SARS-CoV-2), which caused the COVID-19 pandemic and now is endemic across the globe, encode a papain-like protease (PLpro), which is part of the Nsp3 protein embedded into viral replication organelles (also called double-membrane vesicles (DMVs)). PLpro is essential for virus replication and dissemination in the host, and thus a major target for antiviral drug development^9–12^. PLpro participates in viral polyprotein cleavage and also has deubiquitinating (DUB) and deISGylating activities^13^, the two latter functions dysregulating various innate immune and inflammatory pathways^7,12,14–16^. However, the physiological role of coronaviral deISGylating activity in immune escape and viral pathogenesis is still elusive, largely due to the difficulty to specifically ablate PLpro’s deISGylating activity in the context of a recombinant virus. Moreover, coronaviruses such as SARS-CoV-2 are known to alter various host metabolic processes, contributing to immunopathology and severe COVID-19 outcomes^17^; however, whether PLpro-induced deISGylation of host proteins contributes to coronaviral reprogramming of metabolic processes is unknown.

Here, we engineered a recombinant SARS-CoV-2 encoding a mutant Nsp3 that is impaired in PLpro deISGylating activity. Infection studies in human cells and in mice showed that this mutant recombinant virus is attenuated due to heightened innate immune antiviral responses. Furthermore, untargeted metabolomics analysis showed that ablated viral deISGylation led to dysregulation of various metabolic pathways during infection. Global ISGylome proteomics profiling, combined with molecular characterization studies, identified key innate immune and metabolic proteins directly targeted by PLpro, unveiling the physiological role and cellular target landscape of SARS-CoV-2-induced deISGylation at the molecular level.

## RESULTS

### A recombinant SARS-CoV-2 impaired in PLpro deISGylation activity is attenuated and elicits heightened innate immune responses

Coronaviral PLpro binds ISG15 and di-ubiquitin (di-Ub) through a bipartite interface: a proximal ‘site 1’ that recognizes Ub-1 or the C-terminal ubiquitin-like (C-UBL) domain of ISG15 through the PLpro zinc-binding “Fingers” and catalytic “Palm” domains, and a distal ‘site 2’ that binds Ub-2 or the N-terminal UBL (N-UBL) of ISG15 through the PLpro “Thumb” domain^13^ (**Figures 1A and S1A**). Comparative structural analyses of these interfaces have facilitated the design of separation-of-function mutations, yielding Ub-binding-deficient PLpro/PLP recombinant mutants with impaired deubiquitinating (DUB) activity for mouse hepatitis virus (MHV), MERS-CoV, and SARS-CoV-2^18–20^. However, attempts to generate a viable recombinant coronavirus with selectively ablated ISG15 binding of PLpro/PLP have thus far been unsuccessful.

**Figure 1.**
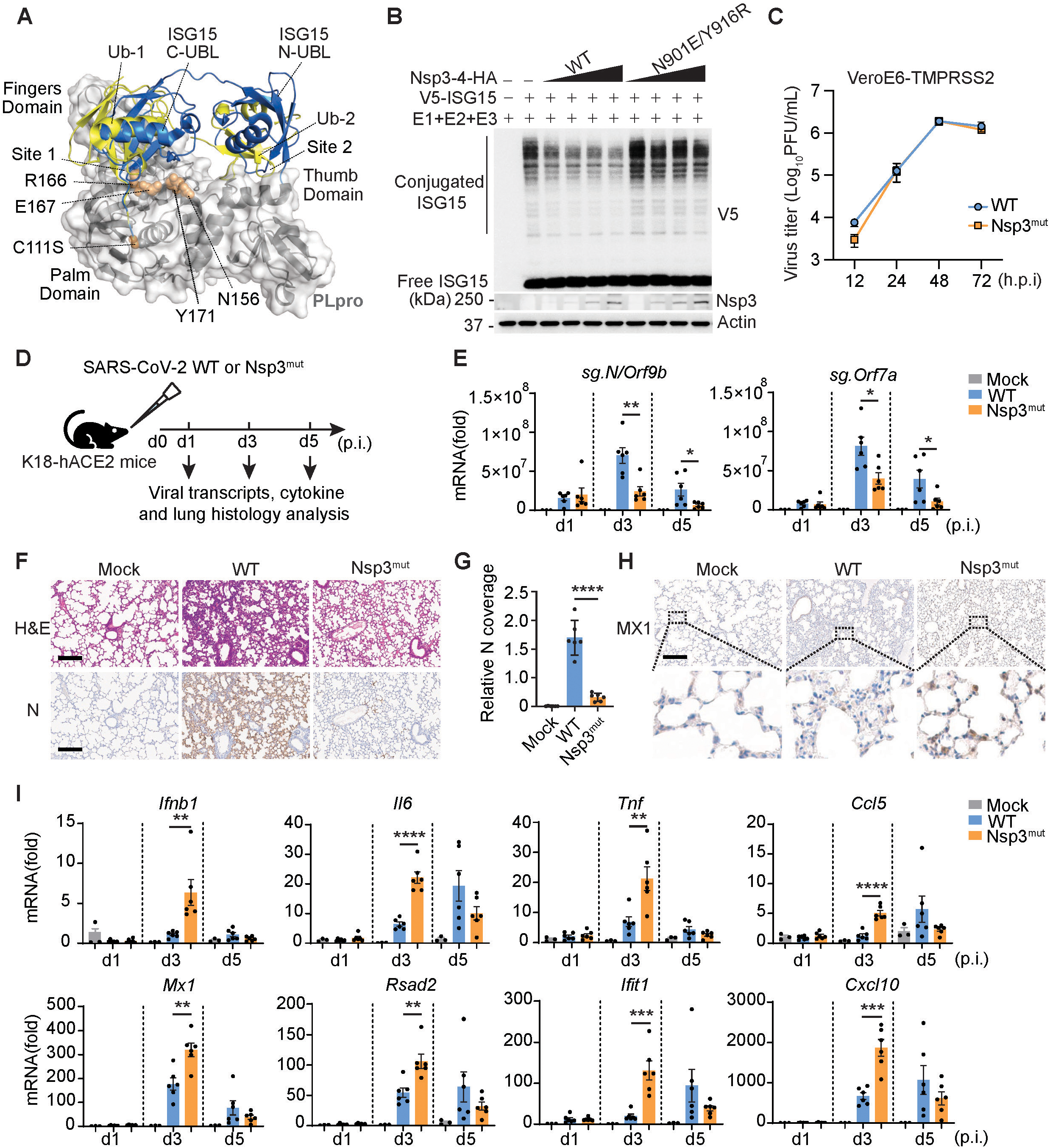
Enhanced innate immune activation and attenuated replication capacity of a recombinant SARS-CoV-2 impaired in PLpro deISGylation activity. (**A**) Superimposed crystal structures of SARS-CoV-2 PLpro catalytic mutants (grey cartoon with white surface, PDB: 7UV5)^71^ complexed with di-Ub (yellow cartoon, PDB: 7UV5)^71^ and ISG15 (blue cartoon, PDB: 7RBS)^71^. While ISG15 and di-Ub occupy both sites 1 and 2 on PLpro, ISG15 makes additional contacts with PLpro site 1 through N165 and Y171 on α6 and α7 helices, respectively. (**B**) Global ISGylation in transiently transfected HEK293T cells that were co-transfected for 24 h with either empty vector (−) or HA-tagged SARS-CoV-2 Nsp3-4 WT or mutant together with V5-ISG15 and ISGylation machinery components (E1, E2 and E3), determined by IB with anti-V5. Whole cell lysates (WCLs) were further immunoblotted with anti-Nsp3 and anti-Actin (loading control). (**C**) Viral titers of SARS-CoV-2 WT and Nsp3^mut^ in VeroE6-TMPRSS2 cells at the indicated times post-infection, determined by plaque assay. PFU, plaque-forming units. (**D**) Schematic of the *in vivo* studies using K18-hACE2 mice that were either mock-treated or infected intranasally with 1,000 PFU of SARS-CoV-2 WT or Nsp3^mut^ (N = 3 for mock; N = 6 for WT or Nsp3^mut^). On day 1, 3, and 5 post-infection (p.i.), lung tissues were collected and subjected to viral transcript and cytokine analysis by RT-qPCR as well as to histological analysis. (**E**) RT-qPCR analysis of viral subgenomic RNAs (sg.) for *N/Orf9b* and *Orf7a* in the lungs of mock-treated or SARS-CoV-2 (WT or mutant)-infected K18-hACE2 mice at the indicated times. (**F**) Hematoxylin and eosin (H&E) staining and immunohistochemistry (IHC) for SARS-CoV-2 nucleocapsid (N) protein in lung sections from K18-hACE2 mice that were mock-treated or infected with SARS-CoV-2 WT or Nsp3^mut^ on day 3 p.i.. Scale bars, 200 μM. (**G**) Relative abundance of viral N protein from the experiment in (F), quantified as described in the Methods. (**H**) MX1 protein expression in lung sections from K18-hACE2 mice that were mock-treated or infected with SARS-CoV-2 WT or Nsp3^mut^, determined by IHC analysis on day 3 p.i. Scale bar, 200 μM. (**I**) RT-qPCR analysis of the indicated cytokine, chemokine, or ISG transcripts in the lungs of K18-hACE2 mice that were mock-treated or infected with SARS-CoV-2 WT or Nsp3^mut^ for the indicated times. Data are representative of at least two independent experiments [mean ± SD of N = 3 (**C**) or N = 6 (**E**, **G** and **I**) biological replicates]. *P < 0.05, **P < 0.01, ***P < 0.001, ****P < 0.0001 (unpaired Student’s *t* test) h.p.i., hours post-infection. d.p.i., days post-infection.

In SARS-CoV-2 PLpro, structural overlays of ISG15 C-UBL and Ub-1 at the site 1 interface highlight distinct binding poses that suggest the feasibility of selectively disrupting ISG15 interaction while preserving Ub binding. While both ISG15 C-UBL and Ub-1 contact residues R166 and E167 at the proximal end of α-helix 7, ISG15 C-UBL is rotated relative to Ub-1 to further interact with the distal end of α-helix 7 (Y171) and the tip of α-helix 6 (N156), distinguishing its interaction from that of Ub^13^ (**Figures 1A and S1A**). This unique binding mode is conserved across human and mouse ISG15 (**Figure S1B**). We thus focused on these site 1 contacts as candidate residues for engineering separation-of-function mutations to specifically ablate PLpro’s deISGylating activity. Using a cell-based assay that measures deISGylation of the caspase activation and recruitment domains (CARDs) of the RNA sensor MDA5, a direct substrate of SARS-CoV-2 PLpro^7^, we found that PLpro mutations targeting the proximal end of α-helix 7 (i.e., R166S/E167R, E167R, and R166K/E167R) abolished deISGylating activity, with comparable effects as seen with the catalytically inactive C111A mutant (**Figure S1C**). However, attempts to rescue viable mutant recombinant viruses bearing these substitutions were unsuccessful, consistent with recent findings^20^.

Given the proximity of α-helix 7 to the catalytic pocket of PLpro, we hypothesized that targeting the distal end of α7 and the tip of α6, specifically residues Y171 and N156 (**Figures 1A, S1A and S1B**), might preserve PLpro’s essential protease activity while impairing its ISG15-binding ability. N156E and Y171R single mutations moderately reduced MDA5-2CARD deISGylation (**Figure S1C**), whereas the combination (N156E/Y171R) resulted in a greater loss of deISGylating activity against both MDA5-2CARD and full-length MDA5 (**Figures S1C and S1D**). Of note, whereas several of the PLpro mutants exhibited reduced protein stability, the protein expression of the N156E/Y171R double mutant was comparable to that of wild-type (WT) PLpro (**Figure S1C**). In an in vitro cleavage/deconjugating activity assay the PLpro N156E/Y171R mutant displayed a nearly abolished ability to cleave pro-ISG15 but retained K48-tri-Ub cleavage activity comparable to WT PLpro (**Figures S1E and S1F**). By contrast, the catalytically inactive C111A mutant was unable to cleave pro-ISG15 and tri-Ub, as expected^20^. PLpro is part of Nsp3, the largest protein of SARS-CoV-2, that is anchored to and shapes, in conjugation with Nsp4, DMVs where coronaviruses replicate^21,22^. To validate our separation-of-function PLpro mutant at its physiological localization, we generated a Nsp3-4 fusion construct that produces mature Nsp3 and Nsp4 via PLpro-mediated autoprocessing and promotes DMV formation, as previously described^23^. The N901E/Y916R Nsp3-4 mutant (equivalent to N156E/Y171R in the isolated PLpro context) showed a nearly abolished deISGylation activity as compared to WT Nsp3-4 (**Figure 1B**). Notably, the global deISGylation effect of WT Nsp3-4 did not increase dose-dependently (**Figure 1B**), strengthening the concept that DMV localization constrains the substrate specificity of Nsp3-PLpro. Of note, we observed no evident effect of Nsp3-4 WT or mutant on total cellular ubiquitination or K48-linked ubiquitination (**Figure S1G**), suggesting that SARS-CoV-2 PLpro —in the cellular context— is specialized primarily for deISGylation, which is in line with recent findings showing that SARS-CoV-2 PLpro’s DUB activity contributes minimally to viral replication and pathogenesis^20^.

Using reverse genetics, we introduced the PLpro N156E/Y171R double mutation into an ancestral SARS-CoV-2 strain (lineage B1) and successfully rescued a viable mutant virus (hereafter termed Nsp3^mut^) in VeroE6-TMPRSS2 cells. Whole-genome sequencing confirmed the genomic stability of Nsp3^mut^ over at least three passages, with no evidence of reversion or compensatory mutations. Nsp3^mut^ replicated comparably to WT virus and produced comparable levels of dsRNA replication intermediates in VeroE6-TMPRSS2 cells, which are IFN defective (**Figures 1C and S1H**). However, in IFN-competent Caco-2 (intestinal epithelial) cells, Nsp3^mut^ exhibited an approximately 1-log reduction in viral titers early in infection (i.e., at 12 and 24 h.p.i) compared to WT virus (**Figure S1I**), indicating attenuation of the mutant virus. In infection studies using the K18-hACE2 transgenic mouse model (**Figure 1D**), we observed that Nsp3^mut^ exhibited reduced replication in the lungs beginning on day 3 post-infection as compared to the parental virus (**Figure 1E**). Histopathological examination revealed diminished immune cell infiltration and reduced viral antigen (i.e., nucleocapsid (N)) staining in Nsp3^mut^-infected lungs (**Figures 1F and 1G**), consistent with restricted viral replication as compared to WT virus. In accord, Nsp3^mut^ infection elicited enhanced MX1 protein expression (**Figure 1H**), as well as stronger transcript expression of type I IFN (*Ifnb1*), proinflammatory cytokines and chemokines (*Il6*, *Tnf*, *Ccl5*, *Cxcl10*), and ISGs (*Mx1*, *Rsad2*, *Ifit1*) on day 3 post-infection, which subsided by day 5 post infection along with accelerated clearance of Nsp3^mut^ as compared to WT virus (**Figure 1I**).

These results establish N156E/Y171R as a separation-of-function mutation that selectively disrupts PLpro’s deISGylation activity without affecting its protease or DUB function, enabling the rescue of a viable mutant virus (Nsp3^mut^) that elicits heightened innate immune activation and exhibits attenuated replication and accelerated clearance.

### PLpro-mediated deISGylation suppresses distinct innate immune signaling pathways

We extended our *in vivo* findings to human cell infection models to understand the detailed mechanisms of PLpro deISGylation-mediated innate immune suppression. Given the established involvement of the respiratory and intestinal tracts, as well as emerging evidence of CNS involvement in SARS-CoV-2 infection/COVID-19, we examined cells derived from these compartments, specifically the immortalized cell lines—A549^ACE2^ (type II alveolar epithelial), Caco-2 (intestinal epithelial), SVGA^ACE2^ (astrocytic), and HMC3^ACE2^ (microglial) cells as well as primary human bronchial epithelial (HBEpCs) and nasal epithelial (HNEpCs).

We first conducted RNA sequencing (RNA-Seq) analysis in Caco-2 and HBEpCs infected with either WT or Nsp3^mut^ SARS-CoV-2. In both cell types, Nsp3^mut^ infection led to a stronger upregulation of IFN- and ISG-related gene signatures as compared to infection with WT virus (**Figures 2A–2C, and S2A**). Most of the genes upregulated by Nsp3^mut^ (vs. WT virus) were enriched in pathways related to antiviral responses and type I IFN signaling (**Figures S2B and S2C**). Similarly, RT-qPCR analysis showed markedly higher induction of *IFITM1* and *ISG15* mRNA in Nsp3^mut^-infected Caco-2 and HBEpCs compared to WT virus infection (**Figures S3A and S3B**). Infection of Caco-2, A549^ACE2^, SVGA^ACE2^, HBEpC, and HNEpC cells over a 2-day time course revealed that Nsp3^mut^ induced evidently higher expression of ISG transcripts and proteins (e.g., MX1, RSAD2, IFITM1) compared to WT virus, concomitant with reduced levels of viral genomic RNA (*g.Nsp3*) and N protein (**Figures 2D–2G, S3C–S3F**). Together, these results show that loss of PLpro-mediated deISGylation restores efficient antiviral innate immune responses during SARS-CoV-2 infection.

**Figure 2.**
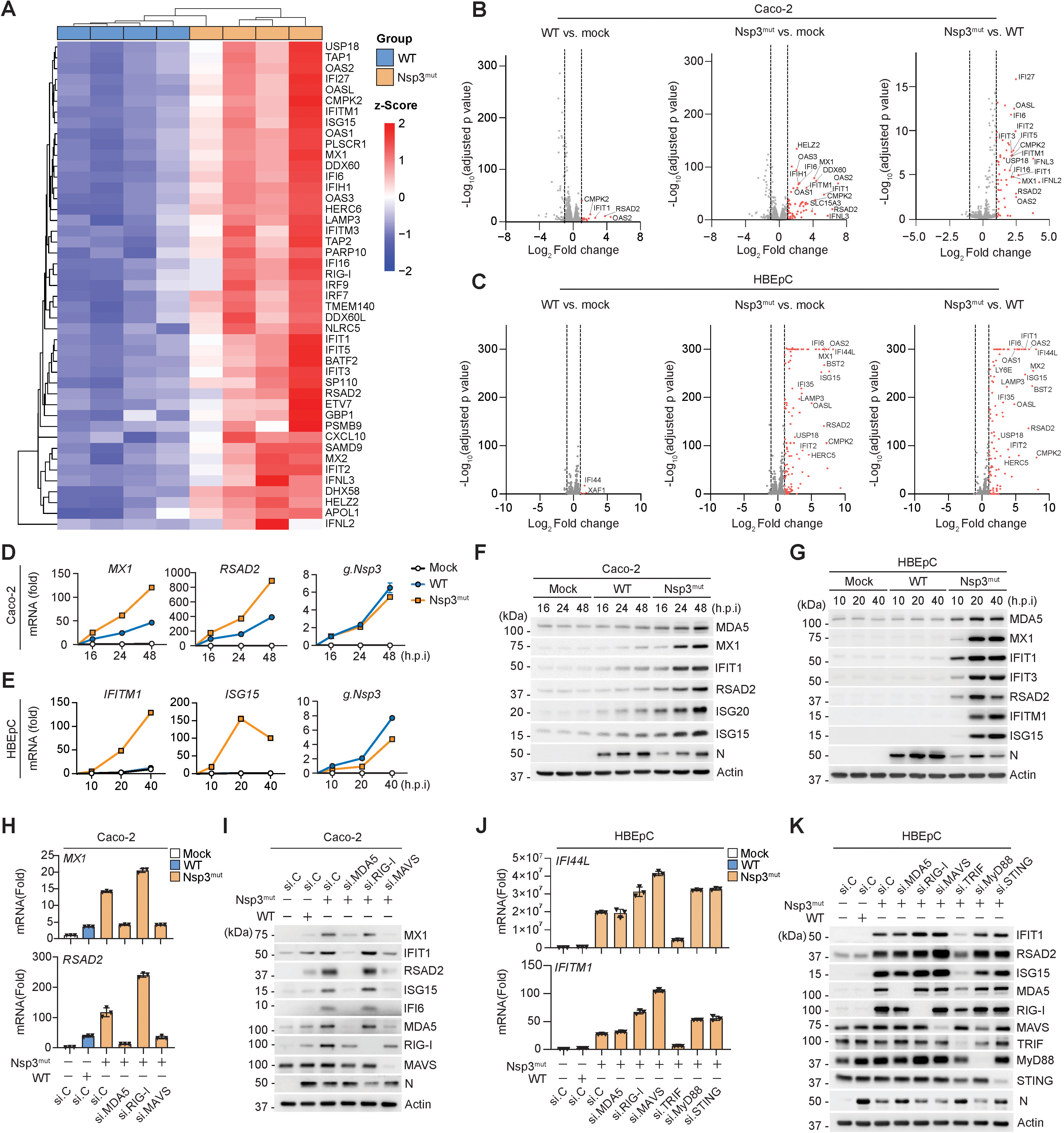
PLpro-mediated deISGylation inhibits MDA5- and TRIF-dependent signaling in a cell type-dependent manner. (**A**) Heatmap showing significantly upregulated genes (fold change > 2, adjusted p-value < 0.05) in Caco-2 cells that were infected for 20 h with SARS-CoV-2 WT or Nsp3^mut^ (each MOI 1), determined by RNA-seq analysis. (**B**—**C**) Volcano plot of differentially expressed genes from the RNA-seq analysis in Caco-2 (**B**) or primary HBEpC (**C**) cells that were either mock-treated or infected with SARS-CoV-2 WT or Nsp3^mut^ for 20 h (see also Methods). Significantly upregulated genes for the indicated comparisons are shown. (**D**—**E**) RT-qPCR analysis of the indicated *ISG* transcripts as well as SARS-CoV-2 genomic RNA (g.) encoding *Nsp3* in Caco-2 (**D**) or primary HBEpC (**E**) cells that were either mock-treated or infected with SARS-CoV-2 WT or Nsp3^mut^ (each MOI 1) for the indicated times. (**F**—**G**) Immunoblot (IB) analysis of the indicated ISGs and viral N protein in the WCLs of Caco-2 (**F**) or HBEpC (**G**) cells that were either mock-treated or infected with SARS-CoV-2 WT or Nsp3^mut^ (each MOI 1) for the indicated times. (**H**) RT-qPCR analysis of *MX1* and *RSAD2* transcripts in Caco-2 cells that were transfected for 48 h with the indicated siRNAs and then either mock-treated or infected with SARS-CoV-2 WT or Nsp3^mut^ for 24 h. si.C, nontargeting control siRNA. (**I**) IB analysis of the indicated ISG and innate signaling proteins from (**H**). (**J**) RT-qPCR analysis of *IFI44L* and *IFITM1* transcripts in primary HBEpCs that were transfected for 48 h with the indicated siRNAs and then either mock-treated or infected with SARS-CoV-2 WT or Nsp3^mut^ (MOI 0.5) for 24 h. (**K**) IB analysis of the indicated ISG and innate signaling proteins from (**J**). Data (**A**—**C**) are from one unbiased RNA-seq screen performed in N = 3 or 4 biological replicates or are representative of at least two (**D**—**K**) independent experiments [mean ± SD of N = 3 biological replicates in (**D**, **E**, **H** and **J**)]. si.C, nontargeting control siRNA.

We next sought to identify the specific innate immune pathways responsible for enhanced sensing of Nsp3^mut^. MDA5 is a major sensor of SARS-CoV-2 (and also other coronaviruses) and is targeted by coronaviral PLpro^7,24^. In addition to MDA5, cGAS and TLR7 have been implicated in SARS-CoV-2-induced IFN responses^25,26^, while TLR1, TLR2, and TLR4 have been reported to recognize viral structural proteins and promote proinflammatory cytokine production^27–30^. In Caco-2 cells, siRNA-mediated knockdown of MDA5 or its downstream adaptor MAVS, but not RIG-I, abolished Nsp3^mut^-induced ISG transcript and protein expression (**Figures 2H and 2I**). Similarly, innate immune activation by Nsp3^mut^ was abolished upon MDA5, but not RIG-I, depletion in SVGA^ACE2^ and HMC3^ACE2^ cells (**Figures S3G–S3I**), strengthening previous findings that PLpro-mediated deISGylation impairs MDA5 signaling as a key immune evasion strategy. Interestingly, in primary HBEpCs, Nsp3^mut^-induced ISG expression was abolished upon TRIF knockdown but remained unaffected by MDA5, RIG-I, MAVS, MyD88 or STING depletion (**Figures 2J and 2K**). In accord, the TRIF-specific blocking peptide Pepinh-TRIF (TFA), abrogated innate immune signaling induced by Nsp3^mut^ in HBEpCs (**Figures S3J and S3K**). Together, these results indicate that PLpro-mediated deISGylation antagonizes at least two major innate sensing pathways —the MDA5-MAVS pathway as well as TRIF signaling— in a cell-type-specific manner.

### Metabolic rewiring via Nsp3-PLpro-mediated deISGylation

SARS-CoV-2 infection is known to profoundly reshape host cellular metabolism to favor viral replication^31–33^. Interestingly, recent evidence linked ISG15 conjugation to distinct metabolic pathways^34^; however, whether viruses manipulate host metabolism through their deISGylation activities is unknown. To determine the role of SARS-CoV-2 PLpro-mediated deISGylation in modulating cellular metabolism, we conducted untargeted metabolomic profiling of primary HBEpCs infected with Nsp3^mut^ or WT virus. Mock-treated cells served as controls (**Figure 3A**). Principal component analysis (PCA) and partial least squares-discriminant analysis (PLS-DA) demonstrated distinct metabolic signatures (**Figures 3B and S4A**). Among the top 20 altered metabolites by WT virus (compared to mock-treated cells) were elevated amino acid derivatives, particularly dipeptides (e.g., Arg-Ala, Ile-Lys, Glu-Gln), which may reflect heightened protein degradation by proteasomal or autophagic pathways under viral stress conditions^31,35^. Furthermore, WT virus infection reduced several lipid species, including phosphatidylethanolamines (PEs), phosphatidylcholines (PCs) and phosphatidylinositols (PIs) (**Figures S4C and S4D**), which is consistent with a previous lipidomic study^36^ and likely due to the extensive membrane rearrangement events occurring during SARS-CoV-2 infection to form DMVs^21,22,37^. By contrast, Nsp3^mut^ infection (compared to mock-treated cells) induced distinct metabolic signatures (**Figures S4E–4G**). Direct comparison of Nsp3^mut^ and WT virus infection revealed a unique metabolic profile associated with impaired PLpro deISGylase activity (**Figure 3C**), and highlighted several key metabolic changes: Nsp3^mut^-infected cells exhibited a significant accumulation of glucose-6-phosphate (G6P), alongside reduced levels of glutathione (GSH), ribose-5-phosphate (R5P), and ribulose-5-phosphate (Ru5P) relative to WT virus-infected cells (**Figures 3D– 3F**). G6P lies at a central metabolic junction between glycolysis and the pentose phosphate pathway (PPP). Since glucose levels remained unchanged (**Figure 3F**), the accumulation of G6P, together with the depletion of downstream PPP intermediates such as R5P and Ru5P, suggests a reduction in PPP flux. Additionally, G6P accumulation may also reflect a downstream block within the glycolytic pathway. This dual bottleneck in glycolysis and PPP may point to inefficient glucose utilization in Nsp3^mut^ infection.

**Figure 3.**
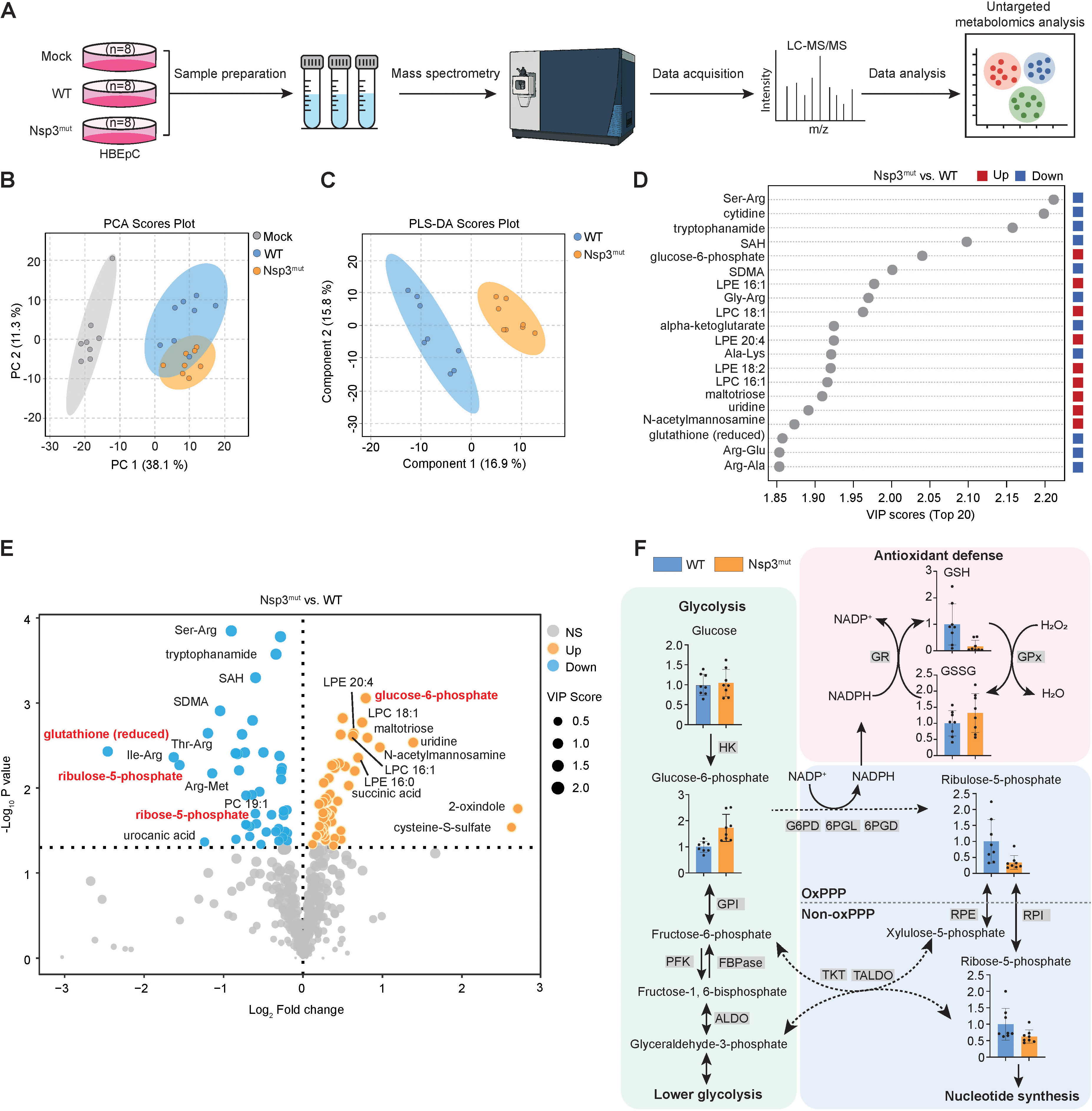
Dysregulation of cellular metabolic processes by Nsp3-PLpro-mediated deISGylation. (**A**) Schematic representation of the workflow for the untargeted metabolomics analysis in primary HBEpCs that were infected with SARS-CoV-2 WT or Nsp3^mut^ (each MOI 1) for 24 h, or that remained uninfected (Mock) (N = 8 biological replicates per condition). Samples were subjected to LC-MS/MS-based metabolite profiling (see Methods for details on sample preparation and metabolomics analysis). (**B**) Principal component analysis (PCA) of global metabolic profiles showing distinct clustering for Mock, WT, and Nsp3^mut^ groups. (**C**) Partial least squares-discriminant analysis (PLS-DA) indicating distinct metabolic signatures of WT vs. Nsp3^mut^ SARS-CoV-2-infected cells. (**D**) Top 20 metabolites contributing to discrimination among groups, ranked by VIP scores from PLS-DA. Metabolites that were significantly upregulated or downregulated in Nsp3^mut^-infected cells (compared to WT SARS-CoV-2-infected cells) are indicated by red and blue boxes, respectively. (**E**) Volcano plot showing significantly altered metabolites in Nsp3^mut^ vs. WT SARS-CoV-2 infection (cutoff: fold change > 1.5, p < 0.05). Notable ISGylation-associated metabolic changes are annotated. (**F**) Schematic summary and bar graphs showing representative metabolites from glycolysis and the pentose phosphate pathway (PPP) that are significantly altered in Nsp3^mut^ infection compared to WT SARS-CoV-2 infection. Data are from one metabolomics profiling screen with N = 8 biological replicates (mean ± SD in (**F**)). Abbreviations: HK, hexokinase; GPI, glucose-6-phosphate isomerase; PFK, phosphofructokinase; FBPase, fructose-1,6-bisphosphatase; ALDO, fructose-bisphosphate aldolase; G6PD, glucose-6-phosphate dehydrogenase; 6PGL, 6-phosphogluconolactonase; 6PGD, 6-phosphogluconate dehydrogenase; RPE, ribulose-5-phosphate epimerase; RPI, ribose-5-phosphate isomerase; TKT, transketolase; TALDO, transaldolase; GR, glutathione reductase; GPx, glutathione peroxidase; GSH, reduced glutathione; GSSG, oxidized glutathione; OxPPP, oxidative branch of the pentose phosphate pathway; Non-oxPPP, non-oxidative branch of the pentose phosphate pathway.

SARS-CoV-2 is known to induce Warburg effect in host cells, favoring aerobic glycolysis over oxidative phosphorylation even in the presence of oxygen. This shift supports rapid ATP generation and supplies biosynthetic precursors for viral replication and assembly^38^. R5P serves as a key precursor for purine and pyrimidine nucleotide synthesis, an essential resource for robust viral RNA replication. The reduced R5P levels in Nsp3^mut^-infected cells (compared to WT virus-infected cells) are expected to limit nucleotide pool availability and thus may contribute to the attenuation of Nsp3^mut^. Furthermore, SARS-CoV-2 infection induces reactive oxygen species (ROS), and GSH is essential for scavenging these reactive species^39^. GSH regeneration depends on NADPH, primarily produced by the oxidative PPP. Thus, reduced GSH levels in Nsp3^mut^-infected cells may result from both elevated oxidative stress and diminished NADPH production due to suppressed PPP activity (**Figure 3F**).

Collectively, these findings show that PLpro-mediated deISGylation critically contributes to SARS-CoV-2-induced reprogramming of several key metabolic processes in infected cells. PLpro’s deISGylation activity contributes to maintaining efficient flux through glycolysis and the PPP, and disruption of this function in the context of Nsp3^mut^ infection leads to impaired biosynthetic and redox capacity, highlighting an important role for viral deISGylation in regulating the metabolic environment necessary for optimal virus replication.

### Global ISGylome and Nsp3-interactome proteomics analyses identify bona fide substrates of PLpro deISGylation

Next, we performed a global ISGylome proteomics analysis to identify substrates of SARS-CoV-2 Nsp3-PLpro (**Figure 4A**). Since WT SARS-CoV-2 encodes a plethora of factors modulating IFN responses and thereby ISG15 levels and ISGylation, making comparative analysis of the ISGylome in Nsp3^mut^ vs. WT virus infection difficult, we expressed in cells WT Nsp3-4 in isolation followed by IFNα treatment to mimic an infected state. Cells expressing empty vector (with or without IFNα stimulation) and cells expressing WT Nsp3-4 without IFNα treatment were included for comparison (**Figure 4A**). Cell lysates were treated with the pan-DUB USP2cc, which removes Ub but not ISG15 from substrates^40^, allowing selective enrichment of ISGylated peptides (**Figure S5A**). Following trypsin digestion and enrichment using anti–K-ε-GG antibody, peptides were analyzed by liquid chromatography-tandem mass spectrometry (LC–MS/MS). This identified a total of 2,736 unique ISGylated peptides across all conditions (**Figure 4B**). Among these, 1,015 ISGylated peptides (corresponding to 578 proteins) were significantly increased (by ≥ 2-fold) upon IFNα treatment but decreased (by ≥ 2-fold) with Nsp3-4 expression, suggesting that these proteins undergo deISGylation —directly or perhaps, indirectly— in the presence of Nsp3-4 (**Figures 4B and S5B**).

**Figure 4.**
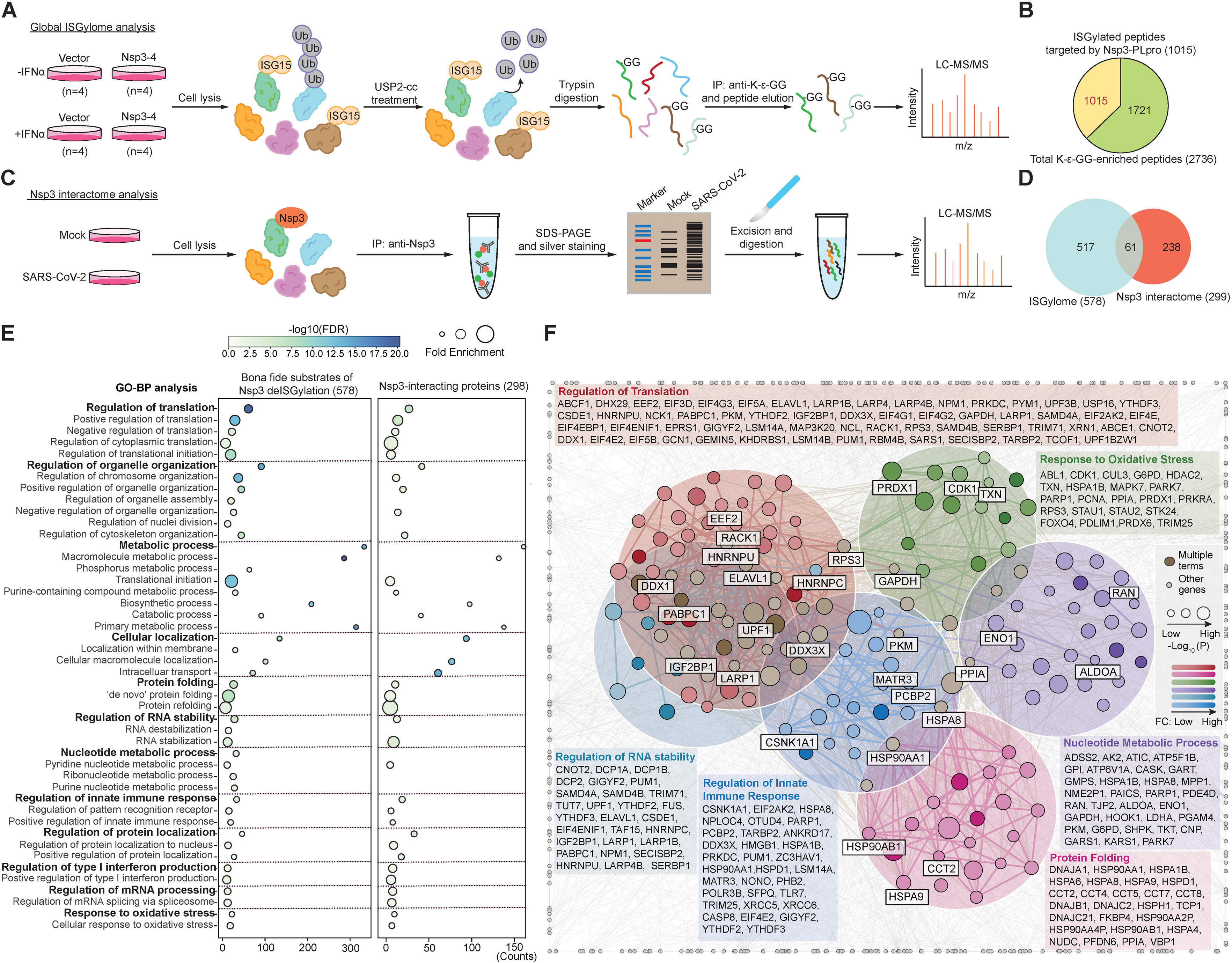
ISGylome and Nsp3-interactome proteomics screens identify bona fide substrates of PLpro deISGylation. (**A**) Schematic representation of the workflow for the global ISGylome proteomics analysis in A549 cells that expressed empty vector or SARS-CoV-2 Nsp3-4 and were either mock-treated (-IFNα) or stimulated with IFNα (+ IFNα) (see also Methods). (**B**) Pie chart showing the total number of ISGylated peptides (2,736) identified from (**A**) and the subset of Nsp3-PLpro–targeted ISGylated peptides (1,015), defined by K^GlyGly^ peptides that had at least a two-fold difference in label free quantification (LFQ) intensity with a p-value < 0.05 in both the comparisons (Vector + IFNα) vs. (Vector - IFNα) and (Vector + IFNα) vs. (Nsp3-4 + IFNα). (**C**) Schematic overview of the workflow of the Nsp3 interactome analysis in Caco-2 cells that were either mock-treated or infected with WT SARS-CoV-2 (MOI 1) for 24 h. Cell lysates were subjected to immunoprecipitation (IP) with anti-Nsp3, followed by LC-MS/MS analysis (see also Methods). (**D**) Venn diagram showing the overlap between the Nsp3-PLpro deISGylation substrates [578 proteins, corresponding to 1015 peptides (**C**)] and the Nsp3-interacting proteins (299 proteins). (**E**) Dot plot of GO enrichment analysis showing functionally convergent biological processes associated with Nsp3-PLpro deISGylation substrates (578) and the Nsp3 interactome. Dot color reflects FDR-adjusted p value, and dot size represents fold enrichment. Only significantly enriched terms (p < 0.05) are displayed. (**F**) Protein-protein interaction network visualization of Nsp3-PLpro deISGylation substrates (578 proteins), generated by Cytoscape. Nodes are color-coded by enriched GO-BP terms: regulation of translation (GO:0006417; red), regulation of RNA stability (GO:0043487; navy blue), regulation of innate immune response (GO:0045088; sky blue), protein folding (GO:0006457; pink), nucleotide metabolic process (GO:0009117; purple), and response to oxidative stress (GO:0006979; green). Enriched proteins are listed under each GO term. Proteins overlapping with the Nsp3 interactome (299 proteins) are highlighted and boxed. Brown nodes represent proteins enriched in multiple terms, and grey nodes represent unenriched proteins. Dot size indicates significance (p value), and color gradient reflects fold change (Vector + IFNα) / (Nsp3-4 + IFNα) in LFQ intensities.

To corroborate direct Nsp3-PLpro deISGylation substrates, we performed an Nsp3 interactome proteomics analysis (**Figure 4C**), allowing us to cross-reference identified interacting proteins with the protein ISGylome dataset. MS analysis of affinity-purified Nsp3 from SARS-CoV-2–infected cells identified 299 interactors (**Figure 4D**). Notably, 61 of these proteins overlapped with the 578 PLpro deISGylation substrates defined by our global ISGylome analysis, indicating that these represent high-confidence PLpro substrates (**Figure 4D**). Comparative Gene Ontology Biological Process (GO-BP) analysis of the two proteomics datasets revealed extensive convergence on several biological processes, including translation regulation, metabolic processes, regulation of RNA stability, regulation of innate immune response, and response to oxidative stress (among others) (**Figure 4E**). Cytoscape network analysis highlighted that the *bona fide* substrates of Nsp3-PLpro’s deISGylation activity converged on six major biological processes (**Figure 4F**).

### Nsp3-PLpro deISGylates key metabolic enzymes in glycolysis and PPP

To investigate the molecular mechanisms by which Nsp3 reprograms host metabolism via PLpro-mediated deISGylation, we looked at the Nsp3-PLpro deISGylation substrates implicated in cellular metabolism identified from our proteomics analysis. This revealed significant enrichment of metabolic enzymes involved in glycolysis, the PPP, amino acid biosynthesis, aminoacyl-tRNA biosynthesis, and carbon metabolism (**Figures 5A and 5B**). Notably, nearly half of these ISGylated metabolic enzymes overlapped with the Nsp3 interactome, and included key regulators of glycolysis and the PPP, such as aldolase A (ALDOA), glyceraldehyde-3-phosphate dehydrogenase (GAPDH), phosphoglycerate dehydrogenase (PHGDH), enolase 1 (ENO1), pyruvate kinase M (PKM), lactate dehydrogenase A (LDHA), and transketolase (TKT) (**Figures 5A and 5C**). These findings suggest that direct deISGylation of these enzymes by PLpro critically contributes to the metabolic reprogramming of infected cells by SARS-CoV-2.

**Figure 5.**
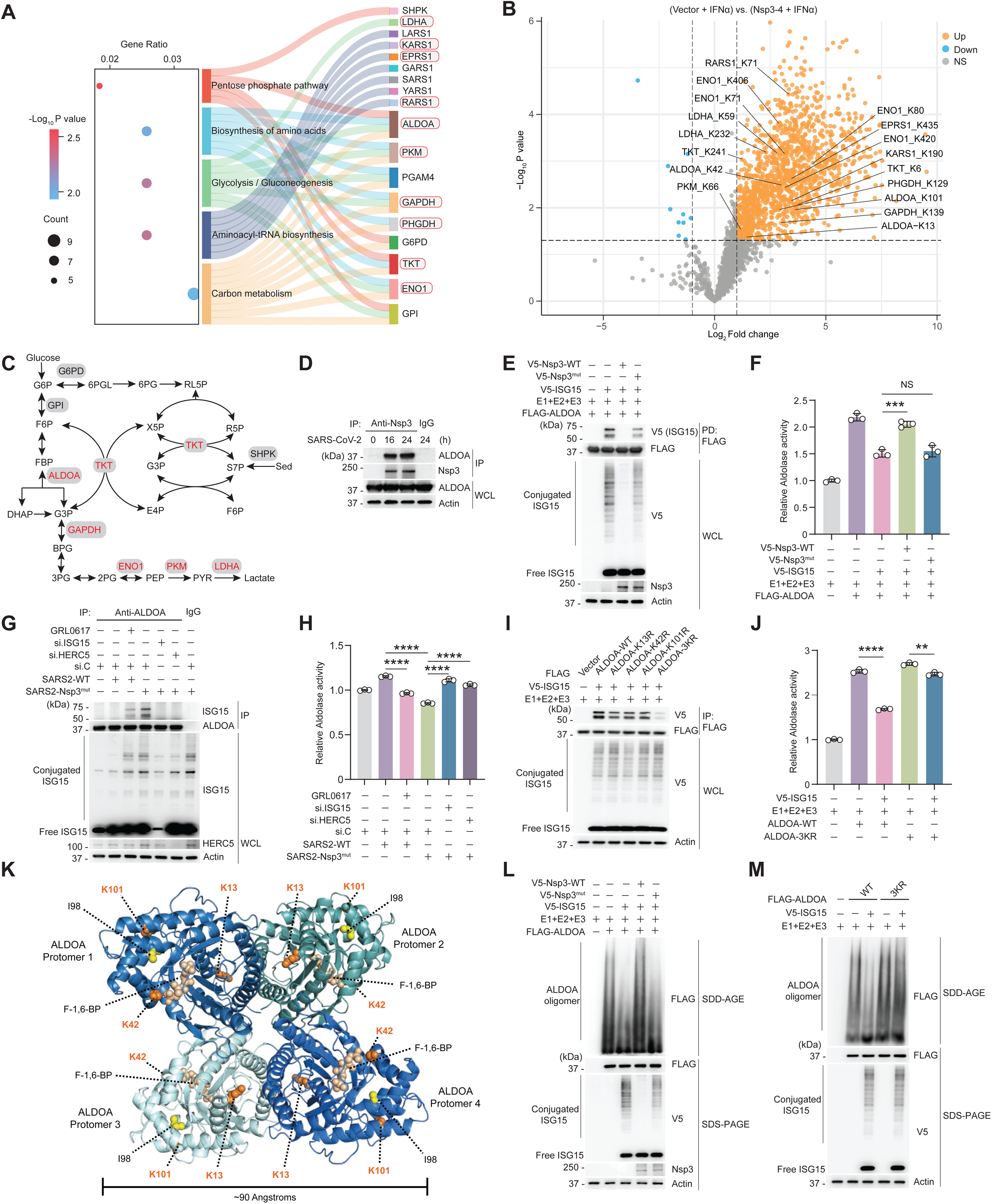
Regulation of key metabolic enzymes in glycolysis and PPP by Nsp3-PLpro deISGylation. (**A**) KEGG pathway analysis showing enriched metabolic pathways among the ISGylated proteins targeted by Nsp3-PLpro (578 proteins). Left, dot plot showing significantly enriched KEGG pathways, ranked by gene ratio and color-coded by –log₁₀(p-value). Dot size represents the number of genes per pathway. Right, Sankey plot illustrating the mapping of ISGylated proteins (right) to their associated KEGG metabolic pathways (center), with pathways grouped by functional categories (left). Proteins that overlap with the Nsp3 interactome are marked with a red box. (**B**) Volcano plot showing the cellular proteins whose ISGylation was significantly reduced by WT Nsp3-4 expression [FC (Vector + IFNα) / (Nsp3-4 + IFNα) >2, *p* < 0.05; orange], with the Nsp3-interacting metabolic enzymes from (**A**) and their specific ISGylation sites (Ks) annotated. (**C**) Schematic diagram illustrating the ISGylated metabolic enzymes involved in glycolysis and the PPP targeted by Nsp3-PLpro deISGylation. The enzymes that overlap with the Nsp3 interactome are highlighted in red font. (**D**) Binding of endogenous ALDOA to Nsp3 in SARS-CoV-2-infected Caco-2 cells at the indicated times post-infection, determined by IP with anti-Nsp3 (IP:Nsp3) and IB with anti-ALDOA and anti-Nsp3. (**E**) ISGylation of FLAG-tagged ALDOA in transiently transfected HEK293T cells that were co-transfected for 24 h with either empty vector (−) or V5-tagged Nsp3 WT or Nsp3^mut^ together with E1, E2, and E3 with or without V5-tagged ISG15, determined by FLAG pulldown (PD:FLAG) and IB with anti-V5 and anti-FLAG. (**F**) Relative aldolase activity in the cell lysates from (**E**). Endogenous aldolase activity from cells co-transfected with empty vector (−) together with E1, E2 and E3 served as control (set to 1). (**G**) ISGylation of endogenous ALDOA in A549^ACE2^ cells that were transfected for 48 h with si.C, si.ISG15 or si.HERC5 and then either mock-treated or infected for 24 h with WT SARS-CoV-2 (with or without GRL0617 pretreatment) or Nsp3^mut^ SARS-CoV-2. (**H**) Relative aldolase activity in the cell lysates from (**G**). (**I**) ISGylation of FLAG-tagged ALDOA WT or mutants in transiently transfected HEK293T cells that were co-transfected for 24 h with either empty vector (−) or V5-tagged ISG15 together with E1, E2, and E3, determined by PD:FLAG and IB with anti-V5 and anti-FLAG. (**J**) Relative aldolase activity in HEK293T cells co-transfected for 24 h with FLAG-tagged ALDOA WT or 3KR together with E1, E2, and E3 with or without V5-tagged ISG15. Endogenous ALDOA activity from cells transfected with empty vector (−) together with E1, E2 and E3 served as control (set to 1). (**K**) Structural visualization of Nsp3-PLpro–targeted ALDOA ISGylation sites in the ALDOA homotetramer (PDB: 4ALD)^44^. ISGylation sites K13, K42 and K101 as well as residue I98 and substrate fructose-1,6-bisphosphate (F-1,6-BP) are highlighted. (**L**) ALDOA oligomerization in transiently transfected HEK293T cells that were co-transfected for 24 h with either empty vector (−) or V5-tagged Nsp3 WT or Nsp3^mut^ together with E1, E2, and E3 with or without V5-tagged ISG15, determined by SDD-AGE and IB with anti-FLAG. WCLs were analyzed by SDS-PAGE and probed by IB with anti-FLAG, anti-V5, anti-Nsp3 and anti-Actin (loading control). (**M**) ALDOA oligomerization in transiently transfected HEK293T cells that were co-transfected for 24 h with either FLAG-ALDOA WT or 3KR together with E1, E2 and E3, with or without V5-tagged ISG15, determined by SDD-AGE and IB with anti-FLAG. WCLs were analyzed by SDS-PAGE and probed by IB with anti-FLAG, anti-V5, and anti-Actin (loading control). Data are representative of at least two independent experiments (**D**—**J**, **L** and **M**). **P < 0.01, ***P < 0.001, ****P < 0.0001 (unpaired Student’s *t* test). NS, not significant; G6PD, glucose-6-phosphate dehydrogenase; GPI, glucose-6-phosphate isomerase; TKT, transketolase; ALDOA, aldolase A; GAPDH, glyceraldehyde-3-phosphate dehydrogenase; ENO 1, enolase 1; PKM, pyruvate kinase M; LDHA, lactate dehydrogenase A; SHPK, sedoheptulokinase; G6P, glucose-6-phosphate; F6P, fructose-6-phosphate; FBP, fructose-1,6-bisphosphate; DHAP, dihydroxyacetone phosphate; G3P, glyceraldehyde-3-phosphate; BPG, 1,3-bisphosphoglycerate; 3PG, 3-phosphoglycerate; 2PG, 2-phosphoglycerate; PEP, phosphoenolpyruvate; PYR, pyruvate; 6PGL, 6-phosphogluconolactone; 6PG, 6-phosphogluconate; RL5P, ribulose-5-phosphate; R5P, ribose-5-phosphate; X5P, xylulose-5-phosphate; S7P, sedoheptulose-7-phosphate; Sed, sedoheptulose; E4P, erythrose-4-phosphate.

To test this hypothesis, we focused on ALDOA, a central regulator of glycolytic flux that influences PPP activity and nucleotide biosynthesis, two processes that are critical for efficient SARS-CoV-2 replication^41–43^. We first validated that endogenous ALDOA is robustly ISGylated in IFN-stimulated *ISG15* (knockout) KO cells reconstituted with WT ISG15 (*ISG15_*GG), but not in cells complemented with the *ISG15_*AA mutant that cannot be conjugated to substrates^7^ (**Figure S6G**). Consistently, IFN treatment inhibited aldolase enzymatic activity in *ISG15*_GG, but not *ISG15*_AA, reconstituted cells, in a dose-dependent manner (**Figure S6H**), indicating that covalent ISG15 conjugation blocks ALDOA activity. Co-Immunoprecipitation (Co-IP) assay confirmed the interaction between ALDOA and Nsp3 when exogenously or endogenously expressed (**Figures S6C and 5D**). Furthermore, ALDOA ISGylation was abolished by co-expression of WT Nsp3, but not Nsp3^mut^, protein (**Figure 5E**). In accord, ISGylation suppressed ALDOA enzymatic activity, which was reversed when WT Nsp3, but not Nsp3^mut^, protein was co-expressed (**Figure 5F**). To determine ALDOA’s ISGylation status during authentic SARS-CoV-2 infection, we analyzed cells infected with WT SARS-CoV-2. No ISGylation of endogenous ALDOA was observed in cells infected with WT SARS-CoV-2; however, pretreatment with the PLpro inhibitor GRL0617 led to prominent ALDOA ISGylation (**Figure 5G**). By contrast, endogenous ALDOA was robustly ISGylated in Nsp3^mut^-infected cells, and ALDOA ISGylation was abolished upon ISG15 or HERC5 silencing, confirming direct ALDOA deISGylation by PLpro during infection (**Figure 5G**). Aldolase enzymatic activity correlated inversely with ALDOA ISGylation, as its activity was increased by WT virus infection, which was reversed upon GRL0617 treatment (**Figure 5H**). Moreover, aldolase enzymatic activity was decreased by Nsp3^mut^ virus infection (as compared to WT virus infection), and this inhibition was reversed by ISG15 or HERC5 silencing (**Figure 5H**). Together, these results indicate that WT SARS-CoV-2, but not the Nsp3^mut^ virus, actively —via PLpro’s enzymatic activity— removes ISGylation from ALDOA, which sustains its efficient enzymatic activity in infected cells.

Our ISGylome MS analysis identified three ISGylation sites in ALDOA: K13, K42, and K101 (**Figure 5B**). Individual mutation of these lysines to arginines modestly reduced ALDOA ISGylation, whereas the combined mutation of all three lysines (3KR) nearly abolished ISGylation (**Figure 5I**). Given that ALDOA consistently displayed two ISGylated bands by western blot, these data suggest that ISGylation of any two of the three lysines may sterically prevent modification of the third lysine. Importantly, unlike ALDOA WT, whose enzymatic activity was markedly inhibited by ISGylation, the activity of the 3KR mutant was much less affected by ISGylation (**Figure 5J**). To further investigate how ISGylation impairs ALDOA enzymatic function, we looked at the identified ISGylation sites in the ALDOA structure^44^. Structural analysis revealed that K13 is situated at the dimer interface between protomers of the ALDOA homotetramer, while ALDOA K42 locates to the α-helix 2 which comprises the active site and is positioned adjacent to the C-terminal tail which caps the active site. Furthermore, K101 is near residue I98, previously implicated in ALDOA catalytic regulation^45^ (**Figure 5K**). Semi-denaturing detergent agarose gel electrophoresis (SDD-AGE) assay showed that ISGylation significantly inhibited ALDOA oligomerization, which was restored by co-expression of WT Nsp3, but not Nsp3^mut^, protein (**Figure 5L**). Consistent with these data, the oligomerization of WT ALDOA, but not of the 3KR mutant, was markedly suppressed by ISGylation (**Figure 5M**).

We also validated G6PD as another bona fide substrate of Nsp3-PLpro. G6PD is the rate-limiting enzyme of the PPP and responsible for generating NADPH for antioxidant defense and R5P for nucleotide synthesis. G6PD ISGylation, induced by co-expression of the ISGylation machinery, was reversed by WT Nsp3, but not by Nsp3^mut^, protein (**Figure S6B**). In line with these results, ISGylation suppressed G6PD enzymatic activity, which was restored by WT Nsp3, but not by Nsp3^mut^, protein (**Figures S6I**). Structural analysis revealed that the identified ISGylation site, K403, is located within the β-sheet of the dimerization domain and makes direct contact with the structural NADP⁺^46^ (**Figure S6J**). Notably, acetylation at K403 has been shown to disrupt dimer formation and abolish enzymatic activity^47^. Given the size of ISG15 (∼54 angstroms in length) (**Figure S6F**), ISGylation at K403 likely sterically hinders G6PD dimerization and NADP⁺ binding, thereby blocking G6PD enzymatic activity.

These data establish ISGylation as a pivotal regulatory mechanism to repress the activity of ALDOA and G6PD by limiting their higher-order assemblies and also show that SARS-CoV-2 PLpro-mediated deISGylation reverses the ISGylation-mediated inhibition of ALDOA and G6PD for creating optimal glycolysis and PPP conditions in the infected cell.

### PLpro-mediated deISGylation of PRDX1 promotes antioxidant responses to prevent viral RNA oxidation

SARS-CoV-2 infection has been widely reported to trigger excessive production of ROS, contributing to oxidative stress, activation of pro-inflammatory responses, tissue damage, and exacerbation of COVID-19 severity^48–51^. Beyond promoting inflammation, elevated ROS levels can directly damage biomolecules, including nucleic acids and proteins. DNA and RNA oxidation, such as the formation of 8-hydroxy-2’-deoxyguanosine (8-OHdG) and 8-hydroxyguanosine (8-OHG), as well as oxidative protein modifications like carbonylation, can have deleterious effects on nucleic acid and protein function and/or stability^52,53^. Given the cytotoxic nature of ROS, how SARS-CoV-2 prevents ROS-induced damage of its viral RNA and proteins remains unknown. We hypothesized that the virus may modulate the activity of key regulators of the host antioxidant response to mitigate oxidative stress and preserve viral integrity.

Our untargeted metabolomics analysis revealed significantly reduced levels of GSH in cells infected with Nsp3^mut^ virus compared to cells infected with WT SARS-CoV-2 (**Figure 3E**), indicating diminished antioxidant capacity and a shift toward oxidative stress upon loss of PLpro’s deISGylation activity. To directly examine oxidative damage, we immunostained for 8-OH(d)G in combination with dsRNA (J2), which marks replicating viral RNA within DMVs. Colocalization immunofluorescence (IF) analysis revealed evidently enhanced oxidation of viral RNA in Nsp3^mut^-infected cells compared to WT virus-infected cells (**Figures 6A and 6B**). Furthermore, global protein carbonylation levels were also markedly elevated in Nsp3^mut^ infection compared to WT SARS-CoV-2 infection (**Figure S6A**). In accord, our gene ontology enrichment analysis of PLpro deISGylation substrates and Nsp3-interacting partners revealed a significant enrichment of pathways in the oxidative stress response (**Figures 4D and 4E**). Among the identified antioxidant substrates, peroxiredoxin 1 (PRDX1) —a key thiol peroxidase that efficiently scavenges hydrogen peroxide (H₂O₂) and organic hydroperoxides— stood out as a top candidate of PLpro substrates. We first validated the ISGylation of endogenous PRDX1 and found that, following type I IFN treatment, a dose-dependent increase in PRDX1 ISGylation was observed in *ISG15*_GG, but not *ISG15*_AA, reconstituted *ISG15* KO cells, with PRDX1 displaying a single ISGylated band (**Figure S6K**). Furthermore, Co-IP assay confirmed the interaction of PRDX1 and Nsp3, both in overexpression conditions and at the endogenous protein level during infection (**Figures S6C and S6D**). Notably, PRDX1 ISGylation was ablated by the co-expression of WT Nsp3, but not by Nsp3^mut^, protein (**Figure 6C**).

**Figure 6.**
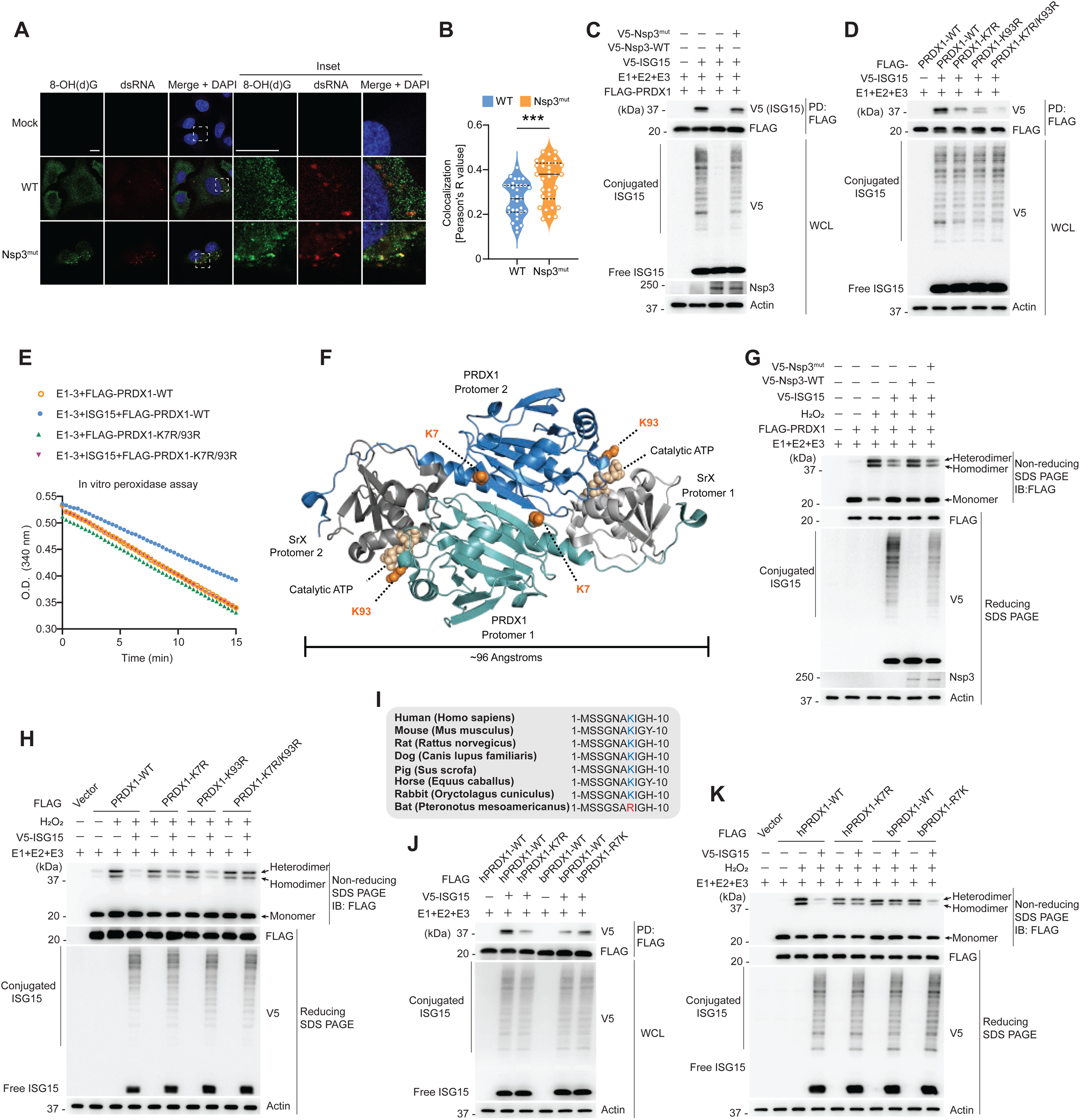
PLpro-mediated deISGylation of PRDX1 enhances antioxidant responses. (**A**) Colocalization of 8-OH(d)G (green) with dsRNA (red) in A549^ACE2^ cells that either remained uninfected (Mock) or that were infected for 24 h with SARS-CoV-2 WT or Nsp3^mut^ (each MOI 2), determined by immunostaining and confocal microscopy analysis. DAPI, nuclei (blue). Scale bar, 10 μm. (**B**) Quantification of 8-OH(d)G-dsRNA colocalization for the data shown in (**A**), determined by Pearson’s correlation coefficient. (**C**) ISGylation of FLAG-tagged PRDX1 in transiently transfected HEK293T cells that were co-transfected for 24 h with either empty vector (−) or V5-tagged Nsp3 WT or Nsp3^mut^ together with E1, E2, and E3 with or without V5-ISG15, determined by PD:FLAG and IB with anti-V5 and anti-FLAG. (**D**) ISGylation of FLAG-tagged PRDX1 WT or mutants in transiently transfected HEK293T that were co-transfected for 24 h with either empty vector (−) or V5-tagged ISG15 together with E1, E2, and E3, determined by PD:FLAG and IB with anti-V5 and anti-FLAG. (**E**) Kinetics of the in vitro peroxidase activity of PRDX1 WT or K7R/K93R purified from mammalian cells and incubated with Trx, TrxR, hydrogen peroxide, and NADPH. Peroxidase activity was assessed by monitoring a decrease in absorbance at 340 nm (A340), which reflects NADPH oxidation. (**F**) Structural visualization of the Nsp3-PLpro–targeted PRDX1 ISGylation sites (K7 and K93) in the PRDX1-SrX complex (PDB: 3HY2)^56^. K7 resides within the dimerization interface, while K93 comprises the shared catalytic active site with SrX that is required for regeneration of PRDX1. (**G**) Dimerization of FLAG-tagged PRDX1 in transiently transfected HEK293T cells that were co-transfected for 24 h with either empty vector (−) or V5-tagged Nsp3 WT or Nsp3^mut^ together with E1, E2, and E3 with or without V5-tagged ISG15. Cells were treated with 100 µM H_2_O_2_ for 5 min before harvesting, and PRDX1 dimerization was assessed by non-reducing SDS-PAGE and IB with anti-FLAG. (**H**) Dimerization of FLAG-tagged PRDX1 WT or mutants in transiently transfected HEK293T cells that were co-transfected for 24 h with either empty vector (−) or V5-tagged ISG15 together with E1, E2, and E3. Cells were treated with 100 µM H_2_O_2_ for 5 min before harvesting, and PRDX1 dimerization was assessed as in (**G**). (**I**) Alignment of the N-terminal PRDX1 protein sequence (amino acids 1–10) from the indicated mammalian species reveals a natural lysine-to-arginine substitution at position 7 (K7R) in PRDX1 from *Pteronotus mesoamericanus*. (**J**) ISGylation of FLAG-tagged human PRDX1 (hPRDX1 WT or K7R) and bat PRDX1 (bPRDX1 WT or R7K) in transiently transfected HEK293T cells that were co-transfected for 24 h with either empty vector (−) or V5-tagged ISG15 together with E1, E2, and E3, determined by PD:FLAG and IB with anti-V5 and anti-FLAG. (**K**) Dimerization of FLAG-tagged hPRDX1 WT or K7R and bPRDX1 WT or R7K in transiently transfected HEK293T cells that were co-transfected for 24 h with either empty vector (−) or V5-tagged ISG15 together with E1, E2, and E3. Cells were treated with 100 µM H_2_O_2_ for 5 min before harvesting, and PRDX1 dimerization was assessed by non-reducing SDS-PAGE and IB with anti-FLAG. Data are representative of at least two (**A**—**E**, **G, H**, **J** and **K**) independent experiments [mean ± SEM of N = 31 cells in (**B**)]. ***P < 0.001 (two-tailed Student’s *t* test with Welch’s correction).

To determine the effect of ISGylation on PRDX1 activity, we performed an in vitro peroxidase assay^54^ using PRDX1 purified from mammalian cells. Non-ISGylated and ISGylated FLAG-tagged PRDX1 were affinity-purified from HEK293T cells expressing ISGylation machinery (i.e., E1, E2 and E3) with or without V5-tagged ISG15. This showed that ISGylated PRDX1 exhibited markedly reduced peroxidase activity as compared to unmodified PRDX1. Furthermore, co-expression of WT Nsp3, but not Nsp3^mut^, protein restored the enzymatic function of ISGylated PRDX1 (**Figure 6E**). Our ISGylome analysis identified K7 and K93 as the specific ISGylation sites in PRDX1 targeted by Nsp3-PLpro (**Figure S5C**). Individual mutation of either site to arginine (K7R or K93R) reduced ISGylation, while the combined mutation (K7R/K93R) almost abolished ISGylation (**Figure 6D**). In line with these results, unlike WT PRDX1, the K7R/K93R mutant retained peroxidase activity in the presence of the ISGylation machinery, strengthening the concept that ISGylation at K7 and K93 suppresses PRDX1 activity (**Figure 6E**).

PRDX1 is a typical 2-Cys peroxiredoxin that functions as a homodimer or higher-order oligomer, with dimerization being triggered by oxidation and required for efficient peroxidase activity and redox cycling^55^. The PRDX1 structure^56^ shows that K7 is located in the N-terminal tail that is involved in dimer formation with the adjacent mirrored protomer, while K93 is close to the shared active site with sulfiredoxin (SrX), which is required for the regeneration of the catalytic C52 of PRDX1 when it becomes hyperoxidized to cysteine-sulfinic acid under oxidative stress (**Figure 6F**). Under normal physiological conditions, PRDX1 undergoes redox cycling in which C52 is oxidized to sulfenic acid upon reacting with H₂O₂, then forms an intermolecular disulfide bond with the resolving cysteine (C173) of the opposite monomer. This disulfide-linked dimer is then reduced back to its active thiol form by thioredoxin (TXN), the primary electron donor in this cycle. Structure modeling of the PRDX1–TXN complex showed that TXN binds at an interface overlapping with that of SrX, suggesting that both enzymes interact with a similar surface around K93 (**Figure S6L**). Thus, ISGylation at K93 is expected to sterically hinder PRDX1 recycling by both SrX and TXN.

To decipher how ISGylation affects PRDX1 dimerization, we performed non-reducing SDS-PAGE and observed that H₂O₂ treatment promoted PRDX1 dimer formation, as expected. By contrast, PRDX1 dimerization was markedly reduced upon ISGylation, and this inhibition was reversed by WT Nsp3, but not by Nsp3^mut^, protein expression (**Figure 6G**). Notably, while the K93R mutant remained sensitive to ISGylation-induced disruption of dimerization, both the K7R and K7R/K93R mutant maintained dimer formation under ISGylation conditions (**Figure 6H**), indicating that reversible ISGylation at specifically K7 is the key regulatory mechanism of PRDX1 dimer formation. Interestingly, protein sequence alignment of PRDX1 across mammalian species revealed that *Pteronotus mesoamericanus*, a bat species known to carry beta coronaviruses^57^, harbors a natural K7R substitution (**Figure 6I**). Compared to human PRDX1 (hPRDX1), PRDX1 from bats (bPRDX1) exhibited significantly reduced ISGylation levels, which were comparable to the hPRDX1-K7R mutant. By contrast, mutation R7 in bPRDX1 to lysine (bPRDX1-R7K) restored ISGylation (**Figure 6J**). Consistent with these data, ISGylation-mediated dimerization inhibition was observed for hPRDX1 and bPRDX1-R7K, but not for hPRDX1-K7R or WT bPRDX1 (**Figure 6K**). Given the role of bats as viral reservoirs, these species likely experience sustained IFN signaling and ISGylation, which may have led to the evolution of the K7R substitution in bPRDX1 and reduced sensitivity to regulation by ISGylation, potentially preserving bPRDK1’s antioxidant function.

Taken together, these results demonstrate that PRDX1 is a key substrate of SARS-CoV-2 Nsp3-PLpro-mediated deISGylation, an event that protects viral RNA within replication organelles from oxidative damage. ISGylation of PRDX1 impairs its dimerization and thereby enzymatic activity, compromising cellular antioxidant capacity during virus infection. The PLpro deISGylating activity of Nsp3 rescues PRDX1 function, highlighting a previously undiscovered viral strategy to mitigate intracellular oxidative stress during infection. Moreover, the evolutionary adaptation of PRDX1 in bats may reflect a host mechanism to preserve redox homeostasis in conditions of high ROS levels induced by virus infection or other high metabolic activity unique to bats (e.g., powered flight).

## DISCUSSION

Our study defines the landscape by which SARS-CoV-2 dysregulates host cellular immunity and co-opts metabolism through PLpro-mediated deISGylation. By generating a separation-of-function mutant recombinant virus (Nsp3^mut^) defective in deISGylation activity, and through the integration of multi-omics analyses and functional characterization, we demonstrate that PLpro’s deISGylating activity is critical for coronaviral fitness, immune evasion, and metabolic rewiring. The Nsp3^mut^ virus elicited markedly elevated type I IFN and ISG responses across multiple cell types and in mouse lungs. The potency of immune activation by Nsp3^mut^ was cell type dependent and mediated by MDA5–MAVS signaling in the majority of cell types tested, which is consistent with previous findings showing that ISGylation is required for MDA5 signaling and that, in turn, SARS-CoV-2 PLpro suppresses ISGylation-mediated MDA5 activation^7,58^. Interestingly, in certain cell types (i.e., HBEpCs), we observed that the ISG induction by Nsp3^mut^ did not depend on MDA5, but instead was mediated by TRIF, the canonical adaptor for TLR3 and TLR4. Recent studies have shown that the SARS-CoV-2 spike protein can interact with and activate TLR4, eliciting inflammatory responses^27,29^. Furthermore, SARS-CoV-2 infection activates TLR3 driving cytokine induction^59^, and lower TLR3 expression levels in peripheral blood has been associated with more severe COVID-19 outcomes^60,61^. TRIF-deficient mice infected with SARS-CoV exhibit impaired type I IFN and ISG responses, elevated viral loads, and exacerbated lung pathology^62^. Our data suggests that in the absence of PLpro’s deISGylating activity, TRIF signaling may become uninhibited to drive IFN responses in certain cell types. However, the molecular mechanisms by which ISGylation/deISGylation regulates TRIF signaling during SARS-CoV-2 infection warrants further investigation. Moreover, the precise PAMP(s) activating TLR-TRIF signaling in this context remains to be elucidated. Taken together, these findings support a model whereby PLpro antagonizes both the MDA5–MAVS and TLR3/4–TRIF pathways through deISGylation in a cell-type, and likely tissue microenvironment, specific manner.

SARS-CoV-2 is known to manipulate various metabolic processes, which can contribute to the severity of disease and infection outcome; however, it is unknown whether protein deISGylation by Nsp3-PLpro contributes to viral dysregulation of metabolism in the infected cell. More broadly, how reversible ISGylation regulates cellular metabolic enzymes at the molecular level has been elusive. Our proteomics analyses identified numerous metabolic enzymes that undergo ISGylation in an inflammatory state (i.e., upon IFNα treatment), and some of these enzymes are direct substrates of PLpro’s deISGylation activity. Untargeted metabolomics analysis showed that loss of Nsp3-PLpro deISGylation activity in the context of recombinant SARS-CoV-2 led to disrupted flux in glycolysis and the PPP. Mechanically, our data revealed that ISGylation of ALDOA occurs at specific lysines that are important for ALDOA oligomerization, and accordingly, ISGylation suppressed ALDOA higher-order assembly formation and its enzymatic activity, while PLpro-mediated deISGylation relieved this inhibitory mechanism. In addition to ALDOA, our proteomics analysis identified that several other core glycolytic and PPP enzymes —such as GAPDH, ENO1, PKM, LDHA, and TKT— are ISGylated upon IFNα treatment and, inversely, deISGylated by SARS-CoV-2 PLpro, although the mechanistic role of such ISGylation events in modulating these enzymes warrants future investigation. Our findings highlight a broader role for ISGylation in fine-tuning central carbon metabolism in response to virus infection (or more broadly, inflammatory stress) and indicate that PLpro-mediated deISGylation promotes efficient viral replication by sustaining glycolytic output.

Our work revealed redox homeostasis as a major metabolic process dysregulated by ISGylation/deISGylation. We identified PRDX1 as a direct ISGylation target whose antioxidant activity is suppressed by ISGylation and, in turn, restored by Nsp3-mediated deISGylation. These findings show that PLpro-mediated deISGylation of PRDX1 limits oxidative stress, which intuitively seems contradictory to published reports showing that SARS-CoV-2 (as well as several other viruses) induce oxidative stress in host cells through excessive ROS production, which can contribute to viral pathogenesis by damaging host molecules^63^. However, how viruses tolerate or actively counteract ROS-induced damage to their own components (i.e., viral RNA and proteins) is poorly understood. Indeed, in addition to PRDX1, other key antioxidant enzymes —including thioredoxin (TXN), which reduces oxidized PRDX1, and the 1-Cys peroxiredoxin PRDX6— were also identified as direct deISGylation substrates of Nsp3-PLpro. These findings indicate that SARS-CoV-2 Nsp3 recruits key antioxidant enzymes to the site of replication, the DMVs, and –through active deISGylation by PLpro– maintains their functional activity for preserving viral protein and RNA integrity under oxidative stress. Whether such spatial control of redox balance represents a broader viral adaptation mechanism to hostile oxidative environments, however, requires further investigation.

Our ISGylome proteomics studies identified a broad set of host proteins, and, through biochemical studies, we have validated several of these (i.e., LDHA, ACLY, CDK1, G3BP1, and DDX1) as direct substrates of Nsp3-PLpro deISGylation (**Figures S6B–S6D**). These proteins are involved in metabolic regulation, cell cycle control, and innate immune responses (among several other processes). While the biological role of ISGylation/deISGylation has not been determined for most of these substrates, previous work has shown that ISGylation of LDHA can reduce its enzymatic activity, thereby influencing lactate production under inflammatory conditions^64^. ACLY, a central enzyme in lipid biosynthesis and acetyl-CoA generation, is required for efficient replication of SARS-CoV-2^65^. CDK1 regulates mitotic progression and has also been implicated in type I IFN production at the translational level^66^, suggesting a potential immunoregulatory role during infection. G3BP1 and DDX1 are multifunctional RNA-binding proteins with well-documented roles in innate antiviral responses. DDX1 contributes to TRIF-dependent sensing of viral dsRNA in dendritic cells^67^, while G3BP1 facilitates the translation of multiple ISG mRNAs and is known to positively regulate antiviral innate immune responses^68,69^. Our proteomics screen identified K76 in G3BP1 as a specific ISGylation site, and K76R mutation led to a near-complete abolishment of G3BP1 ISGylation (**Figure S6E**). Structurally, K76 resides in the β-sheet of the NTF2 domain^70^ of G3BP1, which is located opposite to the substrate-binding groove and near the dimerization interface (**Figure S6F**). Further studies are required to elucidate how ISGylation and Nsp3-mediated deISGylation regulate the functions of G3BP1 and other identified PLpro substrates for modulating diverse host processes and viral replication.

Taken together, our work provides mechanistic insights by which SARS-CoV-2 PLpro rewires host cellular responses, in particular innate immunity and metabolic events, through targeted deISGylation. Our global ISGylome screen identified numerous novel substrates of viral deISGylation, which is expected to catalyze multiple new avenues of research for potential therapeutic antiviral or immunomodulatory intervention through host-directed approaches.

## ACKNOWLEDGEMENTS

We thank Ruofan Cao (Cleveland Clinic FRIC Imaging Core), Belinda Willard (Cleveland Clinic LRI Mass Spectrometry Core), and Judith Drazba (Cleveland Clinic LRI Imaging Core). We are grateful to Susan Baker (Loyola University) for helpful discussions, and to Cindy Chiang for technical support with the manuscript. This study was supported by U.S. National Institutes of Health grants R37 AI087846 (to M.U.G), and in part by RF1AG082211 (to F.C).

## AUTHOR CONTRIBUTIONS

Conceptualization, J.Z., G.L., M.U.G.; Methodology, J.Z., G.L., J.X., K.L., C.M.G., H.Y., Z.S., Y.Z., E.W.; Investigation, J.Z., G.L., J.X., K.L., C.M.G., H.Y., Z.S., Y.Z., E.W. Specifically, G.L. initiated the project, designed and generated the mutant virus, and led the characterization of its innate immune activation mechanisms. J.Z. led the in vivo infection analyses, the metabolomics and ISGylome studies, and performed all validation and mechanistic investigations; Writing – Original Draft, J.Z., G.L., M.U.G.; Funding Acquisition, M.U.G.; Supervision, O.F., S.R.S., F.C., M.U.G.

## STAR METHODS

### KEY RESOURCES TABLE

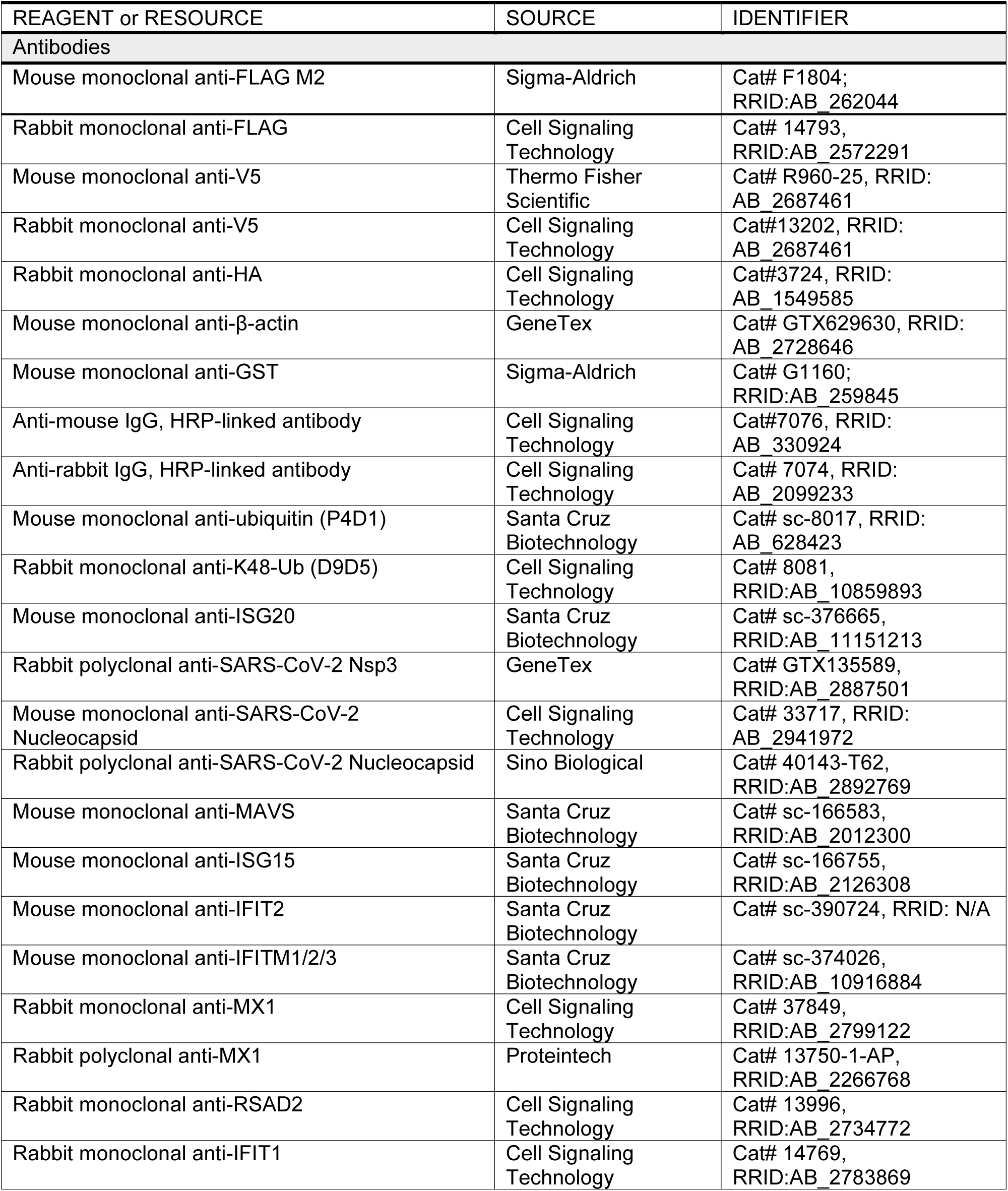
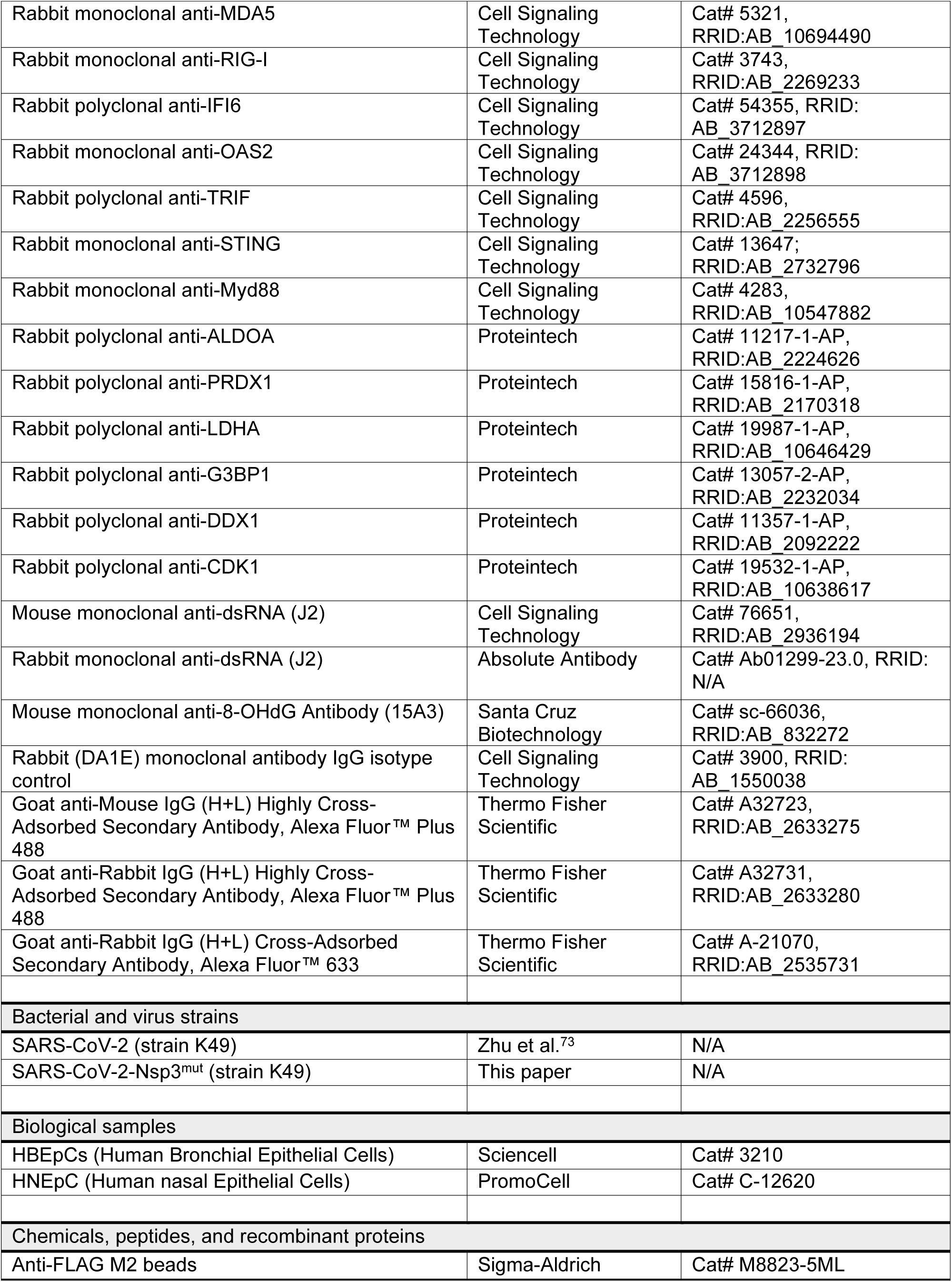
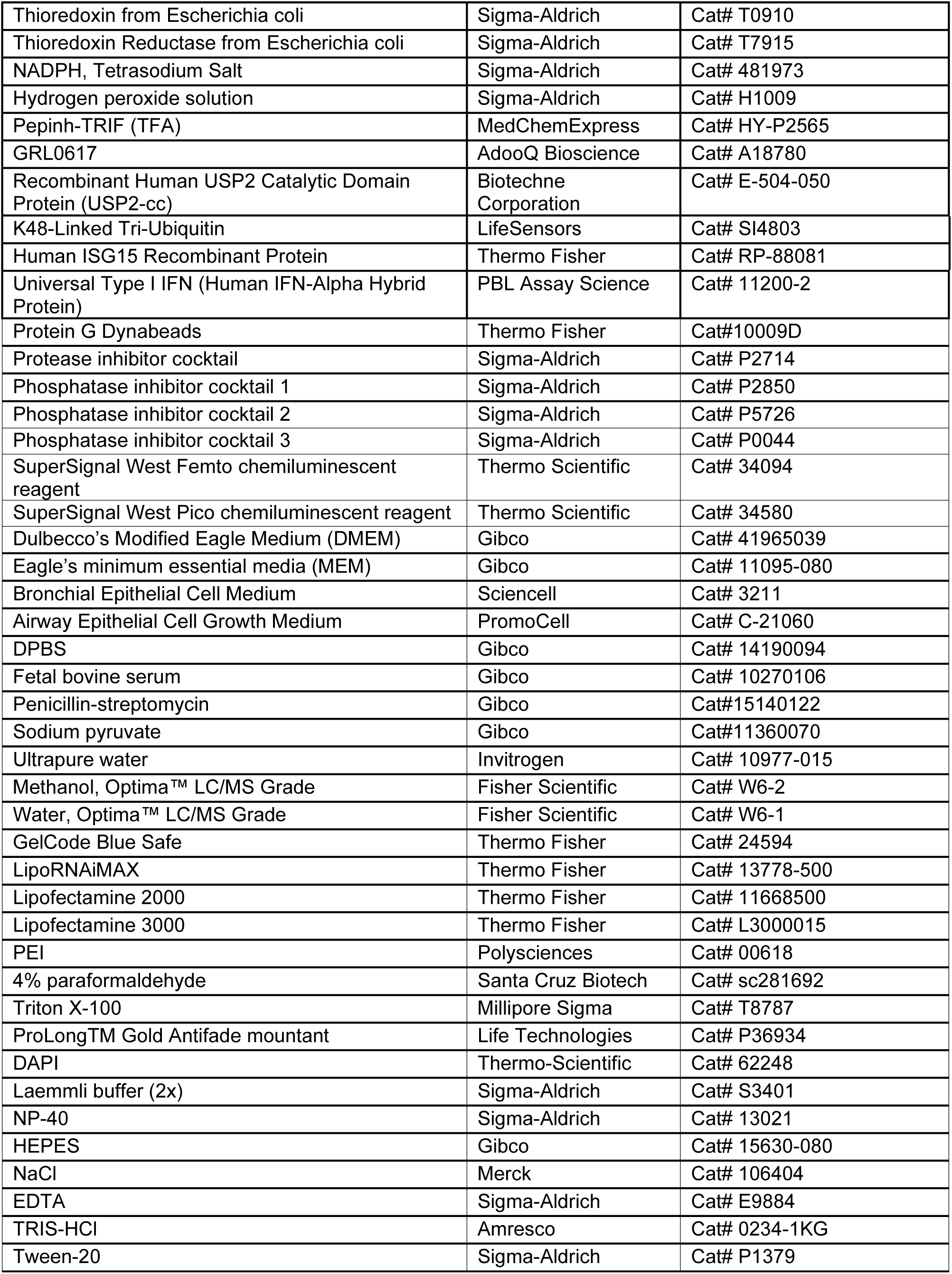
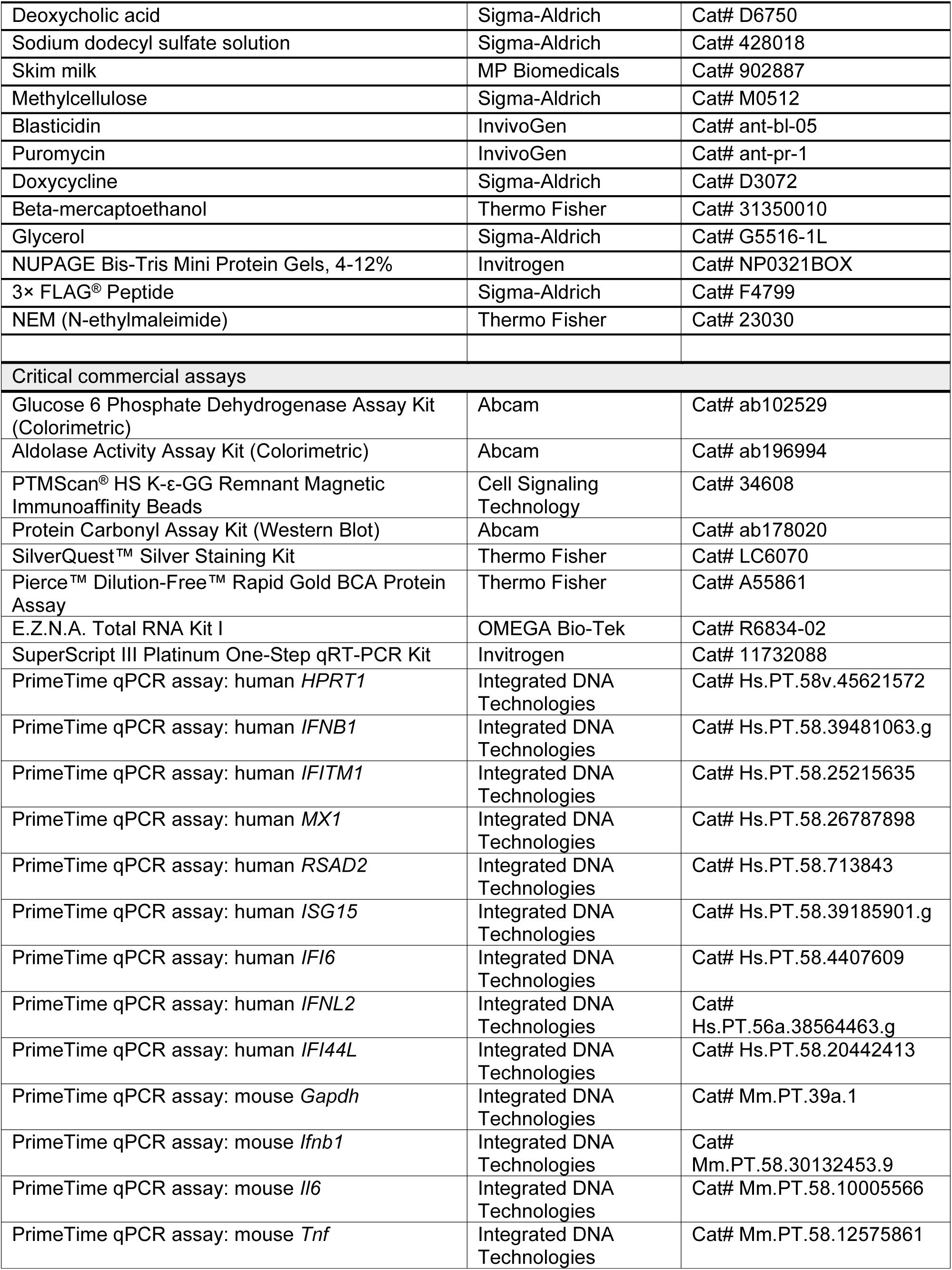
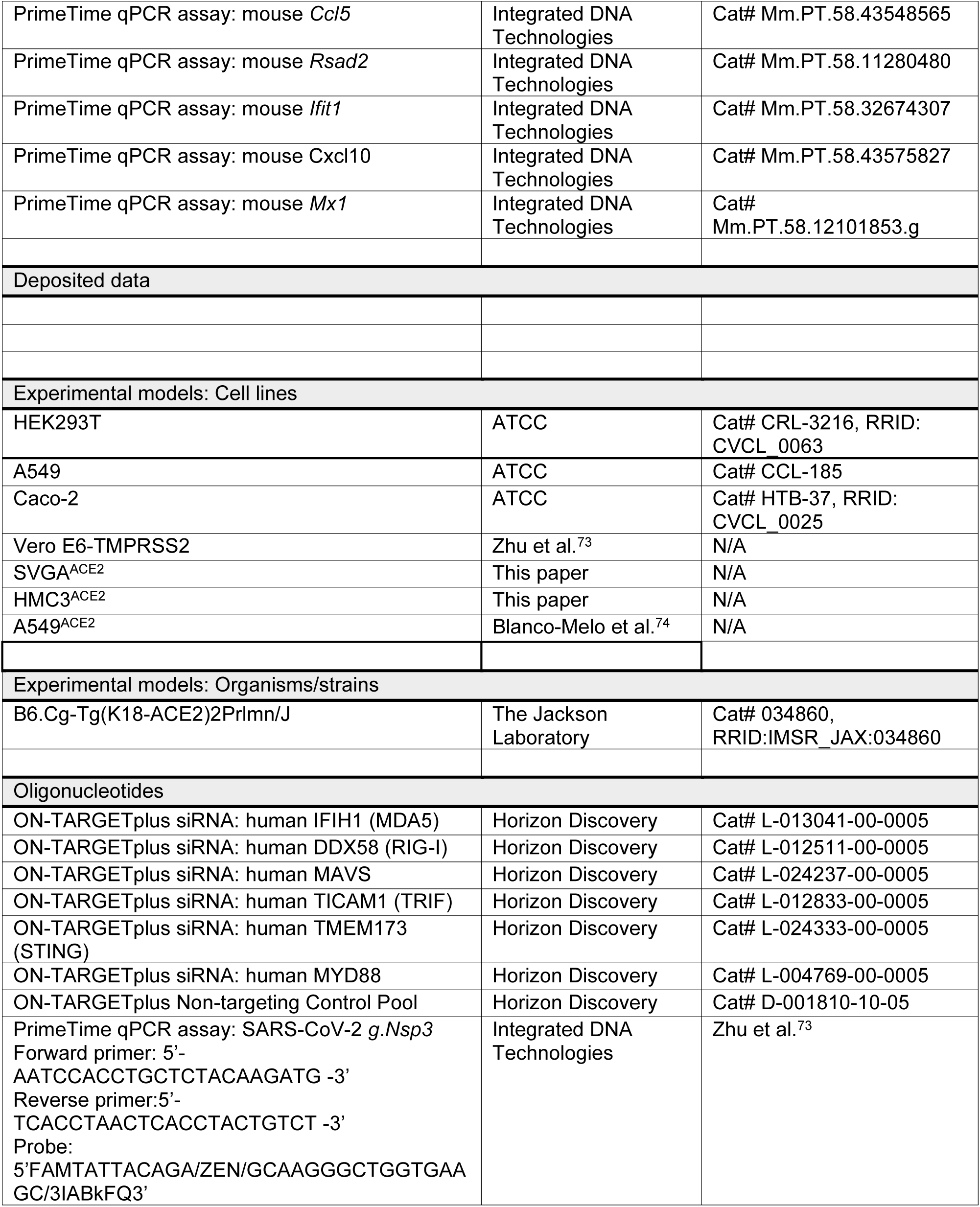
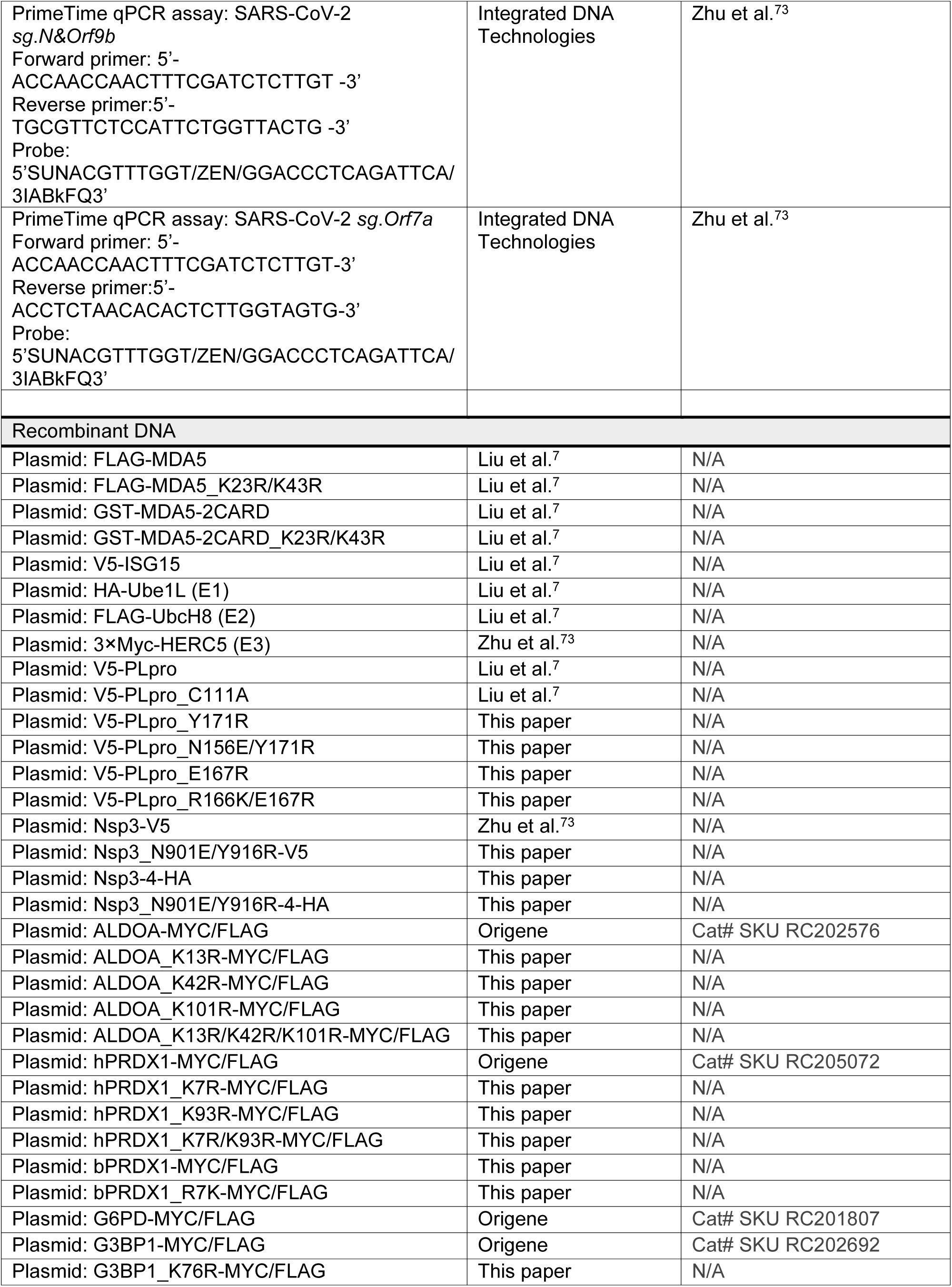
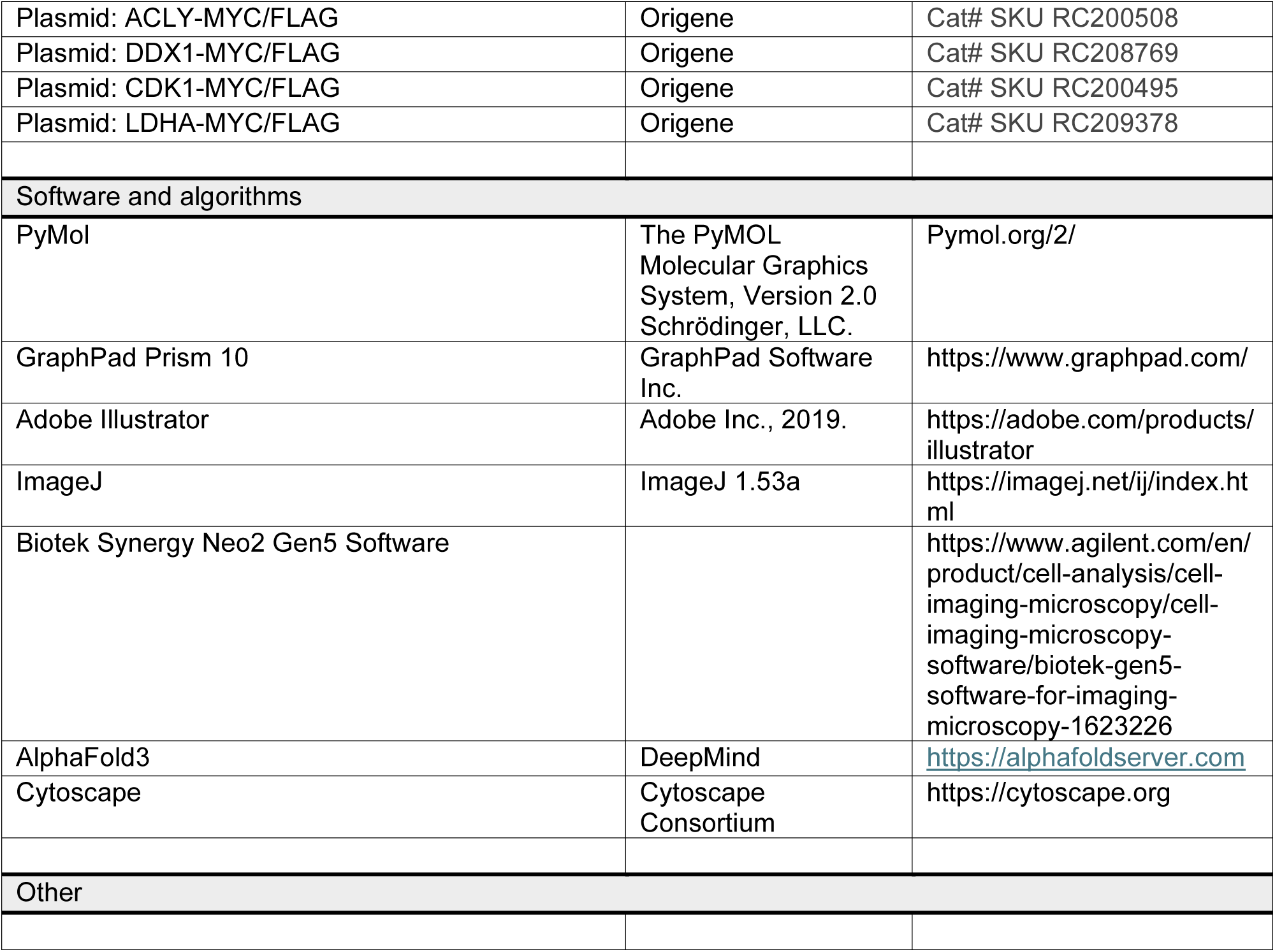

### RESOURCE AVAILABILITY

#### Lead Contact

Further information and requests for resources and reagents should be directed to and will be fulfilled by the Lead Contact, Michaela U. Gack.

#### Materials availability

Reagents and materials generated in this study will be available from the Lead Contact with a completed Materials Transfer Agreement.

#### Data and code availability

The datasets generated during and/or analyzed during the current study are either included in the study and/or available from the corresponding author.

### EXPERIMENTAL MODEL AND SUBJECT DETAILS

#### Cells

HEK293T, A549, A549^ACE2^ were maintained in Dulbecco’s modified Eagle’s medium (DMEM) supplemented with 10% (v:v) fetal bovine serum (FBS), 1 mM sodium pyruvate and 100 U ml^−1^ of penicillin-streptomycin. Vero E6-TMPRSS2 cells were maintained in DMEM supplemented with 10% (vol/vol) FBS, 1 mM sodium pyruvate, 100 U/mL of penicillin–streptomycin, and 40 µg/mL blasticidin. Caco-2 cells were maintained in Minimum Essential Medium (MEM) supplemented with 20% (v:v) FBS, 1 mM sodium pyruvate and 100 U ml^−1^ of penicillin-streptomycin. Human bronchial epithelial cells and Human nasal epithelial cells (HNEpC) were maintained in Airway/Bronchial Epithelial Cell Medium. SVGA^ACE2^ and HMC3^ACE2^ inducibly expressing human ACE2 were generated by lentiviral transduction (pGLVX-TetOne-hACE2) followed by selection with puromycin (1 µg/mL) and maintained in MEM supplemented with 10% (v:v) FBS, 1 mM sodium pyruvate and 100 U ml^−1^ of penicillin-streptomycin.

#### Viruses

SARS-CoV-2 (strain K49) has been described previously^73^ and was propagated in Vero E6-TMPRSS2 cells. Nsp3^mut^ was generated by an Optimized Circular Polymerase Extension Reaction (CPER)-based method^75^ as previously described.

#### Mice

Seven-week-old female K18-hACE2 transgenic mice (B6.Cg-Tg(K18-ACE2)2Prlmn/J) used in this study were obtained from The Jackson Laboratory. Mice were maintained at the Animal Resources Center of the Cleveland Clinic Florida Research and Innovation Center. All mice used in this study were not involved in any other experimental procedure study and were in good health status.

#### Expression constructs

The following plasmids have been described previously^73,75,76^: pcDNA3.1-FLAG-MDA5, pcDNA3.1-FLAG-MDA5_K23R/K43R, GST-MDA5-2CARD, GST-MDA5-2CARD_K23R/K43R, pCAGGS-V5-hISG15, pCAGGS-HA-Ube1L (E1), pFLAG-CMV2-UbcH8 (E2), 3×Myc-HERC5 (E3), pcDNA-V5-Nsp3, pcDNA3.1-V5-PLpro WT and the mutants C111A, R166S/E167R, and N156E. The pcDNA3.1-Nsp3-4-HA construct was generated by fusing Nsp3 and Nsp4 with the natural Nsp3 cleavage linker and cloning the fusion into pcDNA3.1 containing a C-terminal HA tag. Additional mutants were generated by site-directed mutagenesis, including PLpro (Y171R, N156E/Y171R, E167R, and R166K/E167R), Nsp3 (N901E/Y916R, equivalent to N156E/Y171R in the PLpro context), and Nsp3-4 (N901E/Y916R). The pCMV6-Myc/FLAG constructs expressing ALDOA, PRDX1, G6PD, G3BP1, DDX1, CDK1, ACLY, and LDHA were obtained from Origene. The bPRDX1-Myc/FLAG construct was generated by subcloning Pteronotus mesoamericanus PRDX1 cDNA (custom-generated as gBlock, IDT) into the pCMV6-Myc/FLAG vector. Site-directed mutagenesis was used to generate ALDOA (K13R, K42R, K13R/K42R), PRDX1 (K7R, K93R, K7R/K93R), bPRDX1 (R7K), and G3BP1 (K76R) mutants. All constructs were verified by DNA sequencing.

### METHODS DETAILS

#### Transfection and gene silencing

Transient DNA plasmid transfections were performed using linear polyethylenimine (PEI) prepared as 1 mg/mL solution (in 10 mM Tris, pH 6.8), Lipofectamine 2000, or Lipofectamine 3000 following the manufacturer’s instructions.

Transient knockdown in cells was performed using SMARTpool small interfering (si)RNAs as previously described^7^. The specific siRNAs used in our study are described in the Key Resources Table. siRNAs were reverse-transfected into target cells at a final concentration of 80 nM according to the manufacturer’s instructions using Lipofectamine RNAiMAX. At 48 h after transfection, cells were infected with virus or stimulated as indicated in the respective figure legends. Knockdown efficiency was determined by western blotting analysis with the respective antibodies.

#### RNA isolation and RT–qPCR analysis

Total RNA was extracted from cultured cells using the E.Z.N.A. HP Total RNA Kit and from mouse lung tissue using TRIzol Reagent, following the respective manufacturer’s protocols. RT-qPCR were performed using the SuperScript III Platinum One-Step RT-qPCR Kit in combination with PrimeTime qPCR Probe Assays on a QuantStudio 6 Pro Real-Time PCR System (Applied Biosystems). The specific qPCR probes used in our study are described in detail in the Key Resources Table. Relative mRNA expression was normalized to the levels of *HPRT1* or *Gapdh* and expressed relative to the values for control cells using the ΔΔC_t_ method.

#### Co-Immunoprecipitation (Co-IP) and immunoblot analysis

Cells that were transfected as indicated were lysed in 1% Nonidet P-40 buffer (50 mM HEPES pH 7.4, 150 mM NaCl, 1% (v/v) NP-40, 1 mM EDTA, 1× protease inhibitor cocktail) and cleared by centrifugation at 21,000 ×g and 4°C for 15 min. Cell lysates were then subjected to FLAG pulldown (PD) using anti-FLAG (M2) magnetic beads or anti-FLAG M2 agarose beads at 4°C for 4-16 h. For endogenous ALDOA or PRDX1 IP, cell lysates after the indicated treatments were incubated with Protein G Dynabeads conjugated with the indicated antibody at 4°C for 4-16 h. The beads were washed with NP-40 buffer and, afterwards, the proteins were eluted by heating in 1× Laemmli SDS sample buffer at 95°C for 5 min. Protein samples were resolved on Bis–Tris SDS-polyacrylamide gel electrophoresis (PAGE) gels, transferred on to polyvinylidene difluoride (PVDF) membranes, and visualized using the SuperSignal West Pico PLUS or Femto chemiluminescence reagents on an ImageQuant LAS 4000 Chemiluminescent Image Analyzer (General Electric) as previously described^73^.

#### Confocal microscopy

Vero-E6-TMPRSS2 or A549-ACE2 cells grown on 12-mm coverslips were infected with SARS-CoV-2 WT or Nsp3^mut^ at an MOI of 0.5 or 2, respectively. At 24 h post-infection, cells were fixed with 4% (w/v) paraformaldehyde and permeabilized with 0.2% (v/v) Triton-X-100 (in PBS), following protocols previously described^77^. Cells were then blocked with PBS containing 0.05% (v/v) Triton-X-100 and 2% (w/v) BSA for 1 h at room temperature. After washing with PBS, Vero-E6-TMPRSS2 cells were incubated overnight at 4 °C on a rocking platform with mouse anti-dsRNA (J2) antibody diluted in PBS containing 1% (w/v) BSA. A549-ACE2 cells were incubated under the same conditions with rabbit anti-dsRNA (J2) antibody together with mouse anti-8-OHdG antibody. Cells were washed three times with PBS containing 0.25% (v/v) Tween-20 and incubated with Alexa Fluor 488–conjugated secondary antibody for Vero-E6-TMPRSS2 cells, or with Alexa Fluor 488– and Alexa Fluor 633–conjugated secondary antibodies for A549-ACE2 cells. Cells were washed three times with PBS containing 0.25% (v/v) Tween-20, mounted in Prolong Gold Antifade mountant after DAPI staining, and imaged using a Leica Stellaris 8 confocal microscope.

#### Virus titration

SARS-CoV-2 titers were quantified by plaque assay. In brief, monolayers of Vero E6-TMPRSS2 cells were inoculated with ten-fold serial dilutions of virus-containing media. After 2 h, the inoculum was removed, and the cells were washed twice with PBS. The monolayers were then overlaid with 1% colloidal microcrystalline cellulose in MEM supplemented with 2% FBS, 2 mM L-glutamine, 1× non-essential amino acids, 10 mM HEPES, and 100 U/mL penicillin–streptomycin. Plaques were visualized on day 3 by staining with Coomassie Blue. All work in our study relating to SARS-CoV-2 live virus was conducted in the BSL-3 facility of the Cleveland Clinic Florida Research and Innovation Center (CC-FRIC). All work was reviewed and approved by the CC-FRIC Institutional Biosafety Committee in accordance with U.S. National Institutes of Health guidelines.

#### SARS-CoV-2 mouse infections

Seven-week-old K18-hACE2 transgenic mice were intranasally inoculated with 1,000 plaque-forming units (PFU) of either SARS-CoV-2 WT or Nsp3^mut^. Lung tissues were collected at the indicated days post-infection for qPCR, histological, and immunohistochemical analyses. The mouse experiments were carried out in the A/BSL-3 facility of the Cleveland Clinic Florida Research and Innovation Center with the approval of the Institutional Animal Care and Use Committee and in accordance with the National Institutes of Health Guide for the Care and Use of Laboratory Animals.

#### Histology and immunohistochemistry

Mouse lung tissues were fixed in 10% zinc formalin, dehydrated through graded alcohol and xylene baths, embedded in paraffin, sectioned, and stained with hematoxylin and eosin (H&E). Immunohistochemistry (IHC) was performed using the Discovery ULTRA automated staining system (Roche Diagnostics, Indianapolis, IN). For antigen retrieval, tissue sections were treated with Tris/borate/EDTA buffer at pH 8.0–8.5 and heated to 95 °C for 8 min for SARS-CoV-2 nucleocapsid (N) staining or 32 min for MX1 staining. Sections were incubated with primary anti-SARS-CoV-2 N antibody at a dilution of 1:4,000 for 16 min at room temperature, or with primary anti-MX1 antibody at a dilution of 1:800 for 1 h at room temperature. Detection was carried out using the OmniMap anti-Rabbit HRP secondary antibody and visualized with the ChromoMap DAB detection kit. Slides were counterstained with hematoxylin and bluing reagent.

N-protein expression was quantified using ImageJ. Wide-field immunohistochemistry (IHC) images stained for N protein were first converted to RGB color format. Color deconvolution was then applied to separate the DAPI and N protein staining channels. The following RGB vectors were used for the deconvolution: DAPI— [R: 36.24, G: 49.77, B: 60.82], and N protein—[R: 43.32, G: 37.87, B: 24.82]. Binary masks were generated for each channel by applying intensity thresholding: a range of 0– 70 was used for the DAPI channel and 0–90 for the N protein channel. Regions of interest (ROIs) were automatically generated from the binary images using the ‘Create Selection’ function in ImageJ. To quantify N-protein expression, the area of N protein-positive ROIs was divided by the area of DAPI-positive ROIs for each image. These ratios were pooled across all images to obtain the mean and standard deviation of N protein expression levels for each treatment condition.

#### In vitro assay of PLpro DUB and deISGylating activity

V5-tagged PLpro WT or mutant proteins were affinity-purified from transiently transfected HEK293T cells using V5 antibody-conjugated Dynabeads. The immobilized PLpro WT or mutant proteins (∼0.4 μg) were incubated with pro-ISG15 (1.7 μg) and K48-linked tri-ubiquitin (1.5 μg) substrates at room temperature (23°C) in a final reaction volume of 50 μL containing 20 mM Tris (pH 7.5), 100 mM NaCl, and 10 mM DTT. At the indicated times, reactions were stopped by addition of 50 μL 2×Laemmli SDS sample buffer, and cleaved products were analyzed by SDS–PAGE and IB with anti-Ub and anti-ISG15.

#### RNA-seq analysis

Caco-2 cells and HBEpCs were infected for 20 h with SARS-CoV-2 (K49 strain) at an MOI of 1 and 0.5, respectively. Total RNA was extracted using the E.Z.N.A. HP Total RNA Kit, and RNA-seq analysis was performed by AZENTA.

Specifically, RNA samples were quantified using Qubit 3.0 Fluorometer (Life Technologies) and RNA integrity was checked using Agilent TapeStation 4200 (Agilent Technologies). Strand-specific RNA sequencing libraries were prepared by using NEBNext Ultra II Directional RNA Library Prep Kit for Illumina following manufacturer’s instructions (NEB). Briefly, the enriched RNAs were fragmented for 8 min at 94 °C. First strand and second strand cDNAs were subsequently synthesized. The second strand of cDNA was marked by incorporating dUTP during the synthesis. cDNA fragments were adenylated at 3’ends, and indexed adapter was ligated to cDNA fragments. Limited cycle PCR was used for library enrichment. The incorporated dUTP in second strand cDNA quenched the amplification of second strand, which helped to preserve the strand specificity. The sequencing library was validated on the Agilent TapeStation (Agilent Technologies), and quantified by using Qubit 3.0 Fluorometer (ThermoFisher Scientific) as well as by quantitative PCR (KAPA Biosystems). The sequencing libraries were multiplexed and clustered onto a flowcell on the Illumina NovaSeq instrument according to manufacturer’s instructions. The samples were sequenced using a 2×150bp Paired End (PE) configuration. Image analysis and base calling were conducted by NovaSeq Control Software (NCS). Raw sequence data (.bcl files) generated from Illumina NovaSeq was converted into fastq files and de-multiplexed using Illumina bcl2fastq 2.20 software. One mis-match was allowed for index sequence identification.

#### ISGylome proteomics analysis

To identify the global human ISGylome and characterize deISGylation substrates targeted by the SARS-CoV-2 viral deISGylase NSP3, A549 cells (1 × 10⁷ cells per condition) were transfected with either empty vector (HA-vector) or Nsp3-4-HA using Lipofectamine 3000 for 24 h. Cells were subsequently treated with or without IFN-α (500 U/mL) for an additional 24 h. Following treatment, cells were lysed in 1% Nonidet P-40 buffer (50 mM HEPES, pH 7.4; 150 mM NaCl; 1% NP-40; 1 mM EDTA; 1× protease inhibitor cocktail; 10 mM NEM) and 25 μM PLpro inhibitor GRL-0617 to prevent non-specific post-lysis deISGylation by PLpro. Lysates were cleared by centrifugation at 21,000 × g for 15 mins at 4°C. Supernatants were incubated with 1 μM USP2-cc at 37°C for 1 h to specifically remove ubiquitination. Protein concentrations were determined using the BCA assay. For each condition, 2.2 mg of total protein was reduced with dithiothreitol and alkylated with iodacetamide. Proteins were precipitated with cold acetone at -20°C overnight. The precipitated protein was pelleted by centrifugation at 13,000 × g for 10 mins, and the pellets were air-dried. The protein pellets were resuspended in trypsin prepared in 50 mM triethyl ammonium bicarbonate (TEAB) and incubated at 37°C overnight. Digestion was terminated by adding trifluoroacetic acid (TFA) to a final concentration of 0.5–1%. The digests were dried in a SpeedVac and reconstituted in 1.5 mL of 1X HS IAP Bind Buffer and processed with immunoaffinity purification (IAP) using PTMScan® HS K-ε-GG IAP magnetic beads, following the manufacturer’s protocol. The enriched peptides were dried immediately in a SpeedVac, reconstituted in 30 μL 0.1% formic acid, and filtered using a 0.22 µm Millipore Sigma Ultrafree-MC Centrifugal Filter prior to LC-MS/MS analysis. The quantitation of the K^GlyGly^ containing peptides was performed by a label free method that involves aligning the LC chromatograms and comparing the extracted ion chromatograms for each identified peptide and was performed using the program PEAKSOnine. Normalized LFQ intensities were used to perform the binary comparison.

#### SARS-CoV-2 NSP3 interactome during authentic infection

Caco-2 cells were either mock-treated or infected with SARS-CoV-2 (K49 strain) at an of MOI 1 for 24 h. Cell lysates were precleared using Protein G Dynabeads conjugated with IgG isotype control at 4 °C for 1 h, followed by incubation with Protein G Dynabeads conjugated with anti-Nsp3 antibody or an IgG isotype control overnight at 4 °C. The beads were washed four times with RIPA buffer (20 mM Tris-HCl, pH 8.0, 150 mM NaCl, 1% (v/v) NP-40, 1% (w/v) deoxycholic acid, and 0.01% (w/v) SDS), and bound proteins were eluted with 1× Laemmli SDS sample buffer. Samples were resolved by Bis–Tris SDS–PAGE and stained with the SilverQuest™ Silver Staining Kit following the manufacturer’s protocol. Gel lanes were excised and submitted for LC–MS/MS analysis at the Proteomics Core of Cleveland Clinic’s Lerner Research Institute (Ohio).

#### Metabolomics sample preparation

Primary HBEpCs were either mock-treated or infected for 24 h with SARS-CoV-2 WT or Nsp3^mut^ at an MOI of 1. Subsequently, the cells were quenched by methanal. Collected cell samples were stored in methanol at −20 °C prior to metabolite extraction. A biphasic extraction protocol was applied to separate nonpolar lipids and polar metabolites for improved detection coverage based on a previously published method with minor modifications^78^. Initially, cell samples were sonicated in the original methanol solvent for 5 mins, followed by drying using a SpeedVac to remove methanol. To achieve complete cell lysis, the dried samples were homogenized in 225 µL of −20 °C cold methanol containing internal standards, along with 750 µL of methyl tert-butyl ether (MTBE), using a GenoGrinder 2010 (SPEX SamplePrep) for 1 min at 1,350 rpm. The extraction methanol contained the following internal standards for quality control and retention time normalization: sphingosine (d17:1), LPE (17:1), LPC (17:0), MG (17:0/0:0/0:0), DG (12:0/12:0/0:0), PC (12:0/13:0), cholesterol-d_7_, SM (18:1/17:1), ceramide (d18:1/17:0), PE (17:0/17:0), TG (14:0/16:1/14:0)-d_5_, TG (17:0/17:1/17:0)-d_5_, acylcarnitine (18:1)-d3, fatty acid (16:0)-d_3_, MAG (17:0/0:0/0:0), PI (15:0–18:1)-d_7_, PG (17:0/17:0), PS (15:0-18:1)-d_7_, glucosylceramide(d18:1/17:0), mono-sulfo galactosylceramide (d18:1/17:0), and 5-PAHSA-d_9_. The extraction MTBE contained cholesteryl ester 22:1 as an internal standard. The mixture was vortexed for 10 seconds and shaken at 4 °C for 5 mins using an Orbital Mixing Chilling/Heating Plate (Torrey Pines Scientific Instruments). Subsequently, 188 µL of 4 °C cold water was added and vortexed for 20 s to induce phase separation. After centrifugation at 14,000 × g for 2 min, 350 µL of the upper nonpolar phase and two 125 µL aliquots of the lower polar phase were collected and dried. The dried nonpolar layer was reconstituted in 60 µL of methanol/toluene (9:1, v/v) containing an internal standard, [12-[(cyclohexylamino)carbonyl]amino]-dodecanoic acid (CUDA), for reverse-phase (RP) LC-MS/MS analysis. One of the dried polar fractions was reconstituted in 90 µL of acetonitrile/water (4:1, v/v) for hydrophilic interaction liquid chromatography (HILIC) LC-MS/MS analysis, with the following internal standards added: CUDA, caffeine-d_9_, acetylcholine-d_4_, TMAO-d_9_, 1-methylnicotinamide-d_3_, Val-Tyr-Val, betaine-d_9_, acyl carnitine (2:0)-d_3_, N-methyl-histamine-d_3_, l-carnitine-d_3_, butyrobetaine-d_9_, l-glutamine-d_5_, aspartic acid-d_3_, l-arginine-^15^N_2_, cystine-d_4_, asparagine-d_3_, histidine-d_5_, isoleucine-d_10_, leucine-d_10_, methionine-d_8_, ornithine-d_2_, phenylalanine-d_8_, proline-d_7_, threonine-d_5_, tryptohan-d_8_, tyrosine-d_7_, valine-d_8_, spermine-d_8_, glucose-d_7_, fructose-6-phosphate-^13^C_6_, succinic acid-d_4_, taurocholic acid-d_4_, adenosine 5′-monophosphate-^15^N_5_, uridine 5′-monophosphate-^15^N_2_, dopamine-d_4_, taurine-d_4_, uracil-d_2_, biotin-d_4_, N-acetylalanine-d_3_, guanine-^13^C, and adenosine-^13^C_5_. The second dried polar fraction was used for GC-MS analysis following a derivatization procedure. First, carbonyl groups were protected by methoximation. A total of 10 µL of methoxyamine hydrochloride in pyridine (40 mg/mL) was added to the dried sample and incubated at 30 °C for 90 mins. Then, trimethylsilylation was performed by adding 90 µL of N-methyl-N-(trimethylsilyl)trifluoroacetamide (MSTFA) containing C8–C30 fatty acid methyl esters (FAMEs) as internal standards followed by shaking at 37 °C for 30 min. For all three assays including HILIC LC-MS, RPLC-MS and GC-MS analysis, quality control samples were prepared by pooling 10 µL of aliquots from individual cell samples, and method blank samples were prepared using the whole extraction workflow without biological samples.

#### HILIC LC-MS/MS analysis

ipleTOF 6600+ System (SCIEX, Framingham, MA, USA) was coupled with Agilent 1290 Infinity II UHPLC system (Agilent Technologies) was used for HILIC LC-MS/MS analysis with the focus on polar metabolites. HILIC separation was performed on a Waters ACQUITY Premier BEH Amide column (50 mm × 2.1 mm; 1.7 μm) with the VanGuard FIT guard column (Waters Corporation). The column was maintained at 45 °C with flow rate of 0.8 mL/min. For both electrospray ionization (ESI) positive ion mode and negative ion mode, mobile phase A was 100% water with 10 mM ammonium formate and 0.125% formic acid, and solvent B was acetonitrile/water (95/5, v/v) with 10 mM ammonium formate and 0.125% formic acid. Elution gradient was set up as follows: 0 min: 100% B; 0.5 min: 100% B; 1.95 min: 70% B; 2.55 min, 30% B; 3.15 min, 100% B; 3.8 min, 100% B. Sample injection volume was optimized to 5 μL using QC samples. Triple TOF was operated in data-dependent acquisition mode. MS1 parameters were set as follows: mass range: 50-1500 *m/z*; accumulation time: 0.1 secs; period cycle time 0.2501 secs; delay time: 0 secs. Data-dependent MS/MS parameters were set as follows: mass range: 40-1000 *m/z*; accumulation time: 0.02 secs; IDA experiment ion exclusion: for ions smaller than 1000 *m/z*; exclude isotopes within: 4 Da; dynamic background subtract: on; dynamic accumulation: on; auto adjust with mass: on; Q2 transmission window: 30 Da with 49.7% and 100 Da with 50.3%. Mass calibration was performed prior to data acquisition to ensure mass accuracy using APCI Calibration Solution purchased from SCIEX (for positive ion mode: cat. no 4460131; for negative ion mode: cat. no 4460134).

#### RPLC–MS/MS analysis

Thermo Scientific Q Exactive HF-X Hybrid Quadrupole-Orbitrap MS System coupled with Vanquish UHPLC system (Thermo Scientific) was used for RPLC-MS/MS analysis with the focus on nonpolar lipids. RP separation was performed on a Waters ACQUITY Premier BEH C18 column (50 mm × 2.1 mm; 1.7 μm) with the VanGuard FIT guard column (Waters Corporation). The column was maintained at 65 °C with flow rate of 0.8 mL/min. For ESI positive ion mode, mobile phase A was acetonitrile/water (60/40, v/v) with 0.1% formic acid and 10 mM ammonium formate, and mobile phase B was 2-propanol/acetonitrile (90:10, v/v) with 0.1% formic acid and 10 mM ammonium formate. For ESI negative ion mode, mobile phase A was acetonitrile/water (60/40, v/v) with 10 mM ammonium acetate, and mobile phase B was 2-propanol/acetonitrile (90/10, v/v) with 10 mM ammonium acetate. Same elution gradient was used for both ion modes: 0 min: 15% B; 0.75 min: 30% B; 0.975 min: 48% B; 4 min, 82% B; 4.125 min, 99% B; 4.5 min, 99% B; 4.58 min, 15% B; 5.5 min, 15% B. Sample injection volume was optimized to 5 μL using QC samples. The orbitrap MS was operated in data-dependent acquisition mode. The HESI-II ion source conditions were set as follows: spray voltage, 3.6 kV; sheath gas flow rate, 60 arbitrary units; aux gas flow rate, 25 arbitrary units; sweep gas flow rate, 2 arbitrary units; capillary temp, 300 °C; S-lens RF level, 50; Aux gas heater temperature, 370 °C. The following acquisition parameters were used for MS1 analysis: resolution, 60,000, AGC target, 1e6; Maximum IT, 100 ms; scan range 120–1700m/z; spectrum data type, centroid. Data-dependent MS/MS parameters: resolution, 15,000; AGC target, 1e5; maximum IT, 50 ms; loop count, 2; TopN, 2; isolation window, 1.0 m/z; fixed first mass, 60.0m/z; (N)CE/stepped nce, 20, 30, 40; spectrum data type, centroid; minimum AGC target, 8e3; intensity threshold, 1.6e5; exclude isotopes, on; dynamic exclusion, 2.0 s. To improve metabolite annotation, five runs with iterative MS/MS exclusions were performed for both positive and negative electrospray conditions to avoid repeated acquisition of MS/MS spectra.

#### GC-TOF MS analysis

A total of 0.5 μL derivatized sample was injected with 25 s splitless time on an Agilent 6890 GC (Agilent Technologies) using a Restek Rtx-5Sil MS column (30 m × 0.25 mm, 0.25 μm) with 10 m Guard column (10 m × 0.25 mm, 0.25 μm) and 1 mL/min Helium gas flow. The oven temperature was held 50 °C for 1 min, ramped up to 330 °C at 20 °C/min, and held for 5 min. Data were acquired at 70 eV electron ionization at 17 spectra/s from 85 to 500 Da at 1850 V detector voltage on a Leco Pegasus IV time-of-flight mass spectrometer (Leco Corporation). The transfer line temperature was set to 280 °C and ion source temperature was set to 250 °C. Standard metabolites mixtures and blank samples were injected at the beginning of the run and every 10 samples throughout the run for quality control.

#### Metabolomics MS data processing and statistical analysis

Raw GC-TOF MS data were preprocessed by ChromaTOF version 4.50 for baseline subtraction, deconvolution, and peak detection. BinBase was used for metabolite annotation and reporting^79^. Raw LC–MS data were processed using the LC-BinBase database developed at the UC Davis West Coast Metabolomics Center. LC-BinBase uses peak detection and alignment algorithms from MS-DIAL version 4.9. Chromatographic retention times of peaks were normalized using internal standards. LC-BinBase then stores triplets of retention times, accurate mass and mass spectral fragmentation (MS/MS) to define database entries (’bins’). Annotation of the chemical structures of these bins was performed by matching the LC-BinBase MS/MS spectra against all available mass spectral libraries, including the licensed NIST 23 purchased from NIST (https://www.sisweb.com/software/ms/nist.htm), the licensed Agilent METLIN Metabolomics database purchased from Agilent (https://www.agilent.com), the public GNPS database downloaded from the GNPS website (https://external.gnps2.org/gnpslibrary) and the public MassBank of North America database (https://massbank.us). Annotations were conducted by manual verification of peak shapes, retention time prediction matches and MS/MS matches using the entropy similarity algorithm^80^. The detailed parameter setting was as follows: MS1 tolerance, 0.005 Da; MS2 tolerance, 0.01 Da; minimum peak height, 30,000; mass slice width, 0.1 Da; smoothing method, linear weighted moving average; smoothing level, 5 scans; minimum peak width, 10 scans. For metabolites has multiple adducts or detected in two or more assays, values with higher signal-to-noise ratio were kept for quantification. Volcano plot, principal component analysis (PCA), partial least squares discriminant analysis (PLS-DA) were performed using MetaboAnalyst 6.0^81^. Log transformation was applied to improve data normality prior to t-test, auto-scaling were applied prior to PCA and PLS-DA analysis.

#### *In vitro* peroxidase assay

Myc/FLAG-tagged PRDX1 protein was affinity-purified from HEK293T cells using a published FLAG-protein purification protocol^82^ with minor modifications. Briefly, ten 10-cm dishes of HEK293T cells were transfected with Myc/FLAG-tagged PRDX1 WT or mutant, along with E1, E2, and E3 enzymes, together with or without V5-tagged ISG15. After 24 h, cells were washed with PBS and lysed in NP-40 buffer (50 mM HEPES, pH 7.4; 150 mM NaCl; 1% [v/v] NP-40; 1 mM EDTA; protease inhibitor cocktail) on ice for 30 min. Lysates were cleared by centrifugation at 15,000 rpm for 15 min at 4 °C and then incubated with anti-FLAG M2 agarose beads for 4 h at 4 °C. Beads were washed 7–10 times by vigorously vortexing samples at high speed, and bound proteins were eluted with 100 μg/mL 3× FLAG peptide in TBS (10 mM Tris-HCl, 150 mM NaCl, pH 7.4) for 1 h at 4 °C with shaking at 650 rpm. Protein concentrations were determined by BCA assay.

For the *in vitro* PRDX1 peroxidase activity assay, purified PRDX1 WT or mutant (0.5 μM), thioredoxin (Trx, 5 μM), thioredoxin reductase (TrxR, 0.25 μM), and NADPH (200 μM) were mixed in 180 μL of assay buffer (50 mM HEPES-NaOH, pH 7.0; 1 mM EDTA) per reaction in a clear flat-bottom 96-well black plate. The reaction was initiated by adding H₂O₂ at a final concentration of 100 μM. NADPH oxidation was measured by monitoring the decrease in absorbance at 340 nm for 15 min at 30 °C, with readings taken every 20 s using a Synergy Neo2 Hybrid Multi-Mode Reader (BioTek).

#### Semi-denaturing detergent agarose gel electrophoresis (SDD-AGE

ALDOA oligomerization was examined by SDD-AGE using a published protocol^73^ with slight modifications. Briefly, HEK293T cells were transfected for 24 h with Myc/FLAG-tagged ALDOA WT or mutant, along with E1, E2, and E3 enzymes, with or without V5-ISG15. Cells were then lysed in buffer containing 50 mM HEPES (pH 7.4), 150 mM NaCl, 0.5% (v/v) NP-40, 10% (v/v) glycerol, and protease inhibitor cocktail at 4 °C for 20 min. Lysates were cleared by centrifugation at 16,000 × g for 10 min at 4 °C and subsequently incubated on ice for 1 h. Supernatants were then mixed with 4× SDD-AGE buffer [2× Tris/borate/EDTA (TBE), 40% (v/v) glycerol, and 8% (w/v) SDS] to yield a final concentration of 1× (0.5 × TBE, 10% glycerol, and 2% SDS) and incubated at room temperature for 5 min. Samples were resolved on a vertical 1.5% agarose gel prepared in 1× TBE with 0.1% SDS and electrophoresed at 100 V for 30-40 min at 4 °C. Proteins were transferred to a PVDF membrane and detected by immunoblotting with anti-FLAG antibody.

#### Aldolase and G6PD activity assay

The enzymatic activities of ALDOA and G6PD were measured using the Aldolase Activity Assay Kit and the G6PD Activity Assay Kit, respectively, according to the manufacturer’s instructions.

#### PRDX1 dimerization assay

PRDX1 dimerization was assessed using a protocol as previously described^83^. Briefly, HEK293T cells transfected with Myc/FLAG-tagged PRDX1 were either left untreated or treated with 100 µM H₂O₂ for 5 min prior to harvesting. Following removal of the culture supernatant, cells were lysed in 100 μL of alkylation buffer (40 mM HEPES, 50 mM NaCl, 1 mM EGTA, 1% (v/v) CHAPS, protease inhibitor cocktail, 10 μg/mL catalase, and 100 mM NEM). Lysates were then mixed with non-reducing sample buffer (62.5 mM Tris-HCl, pH 7.0, 10% glycerol, 1.5% SDS, and 0.025% bromophenol blue) and subjected to electrophoresis on a 10% Bis-Tris SDS-PAGE gel, detected by immunoblotting with anti-FLAG antibody.

#### Structural Visualization of ISGylation Sites

The coordinate files of experimentally solved structures used to map ISGylation sites were accessed from the RCSB Protein Data Bank^84^. The predicted complex of human TrX and PRDX1 was modeled using AlphaFold3^85^. Figures were rendered using PyMOL (Version 2.5.5).

### QUANTIFICATION AND STATISTICAL ANALYSIS

The data described in our study were analyzed using GraphPad Prism software (version 10). Statistical analyses were performed using a two-tailed Student’s *t*-test (unpaired) unless otherwise stated, and p values of less than 0.05 were considered significant. Significant differences are denoted by *p <0.05, **p <0.01, ***p <0.001 and ****p <0.0001. Pre-specified effect sizes were not assumed, and the number of independent biological replicates (N) is indicated for each figure in the respective legend.

