## Supplementary figures and images for "Dysregulation of innate immunity and cellular metabolism through virus-induced deISGylation"

### Supplementary Figures 1-6

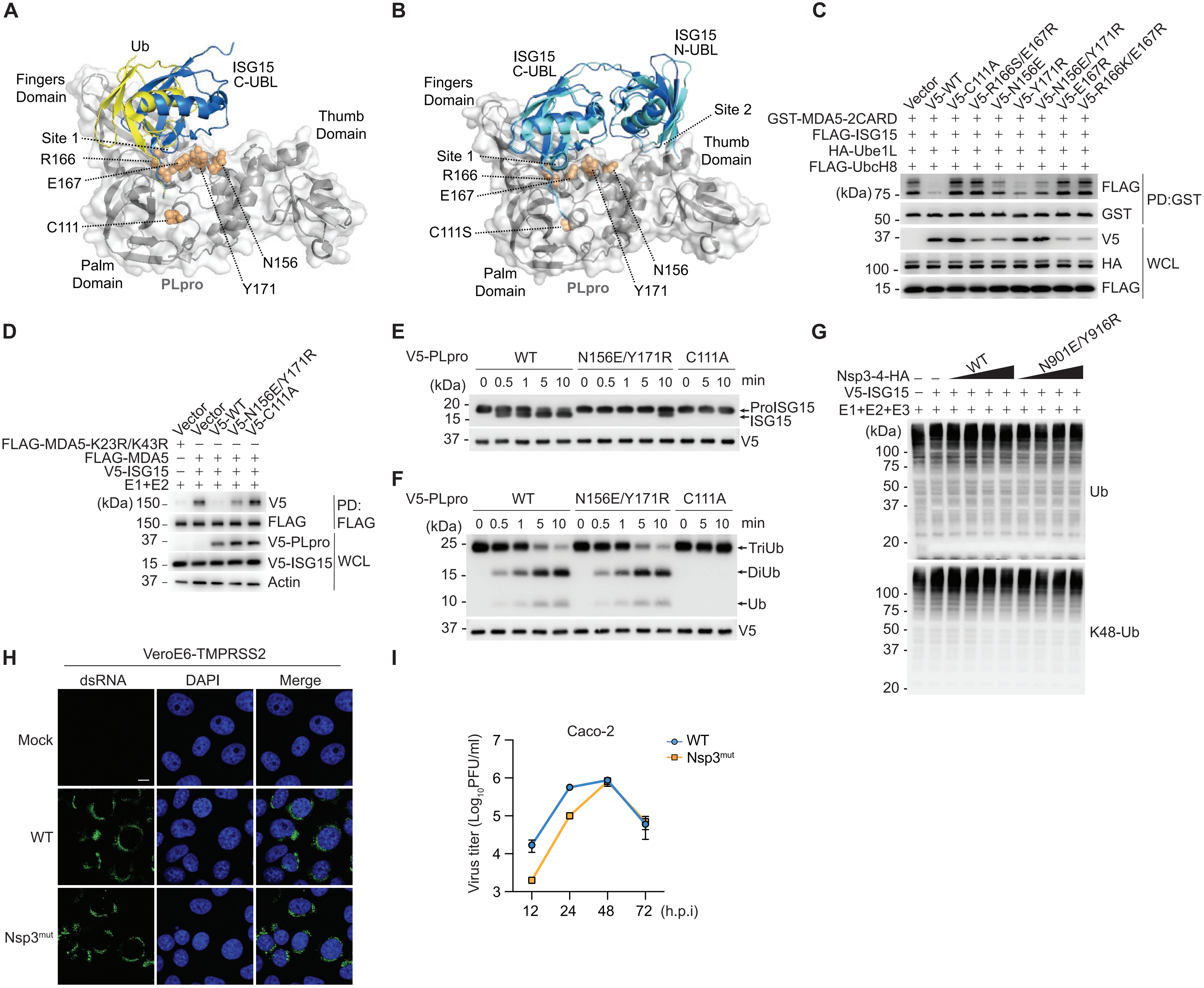

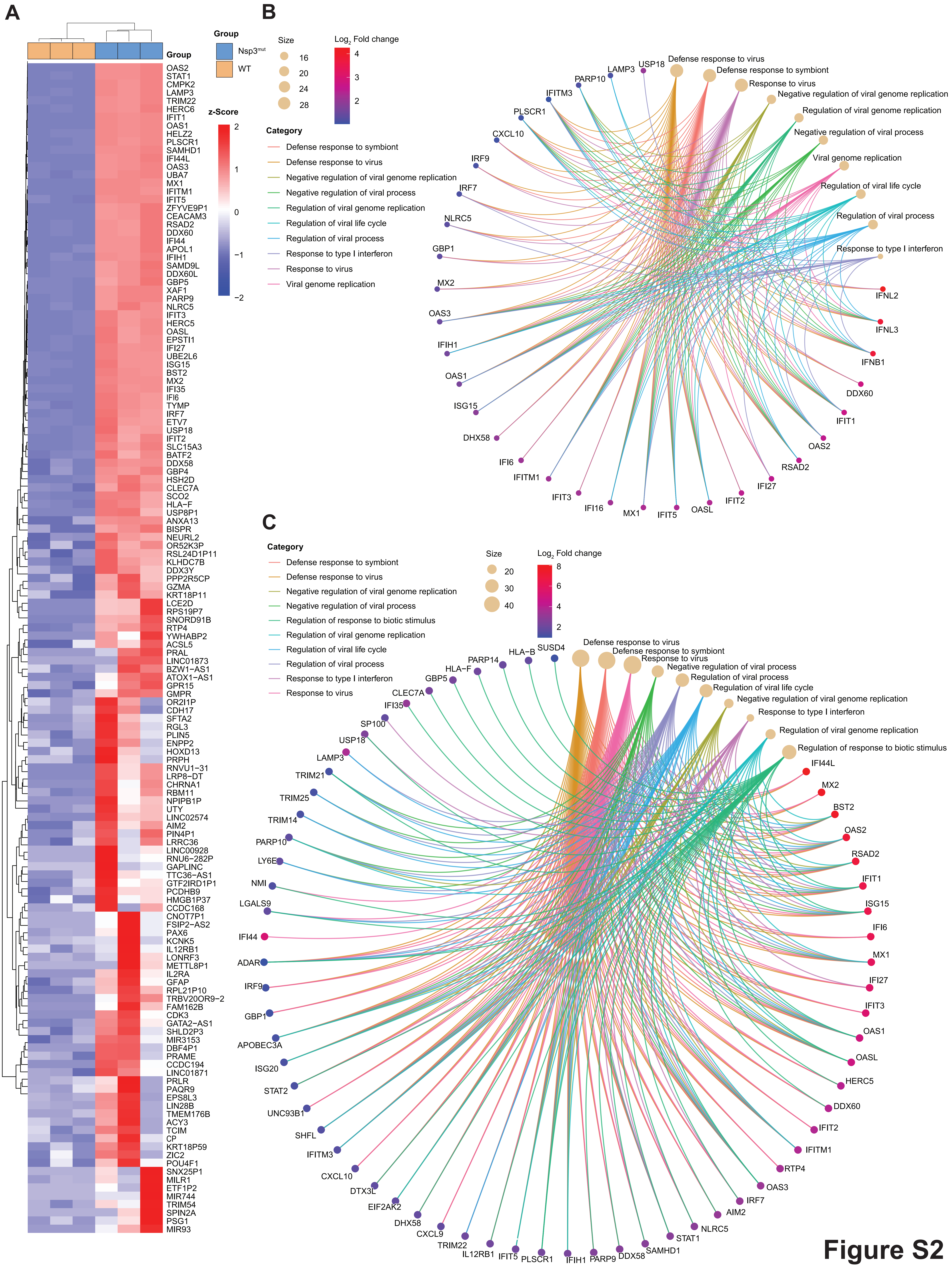

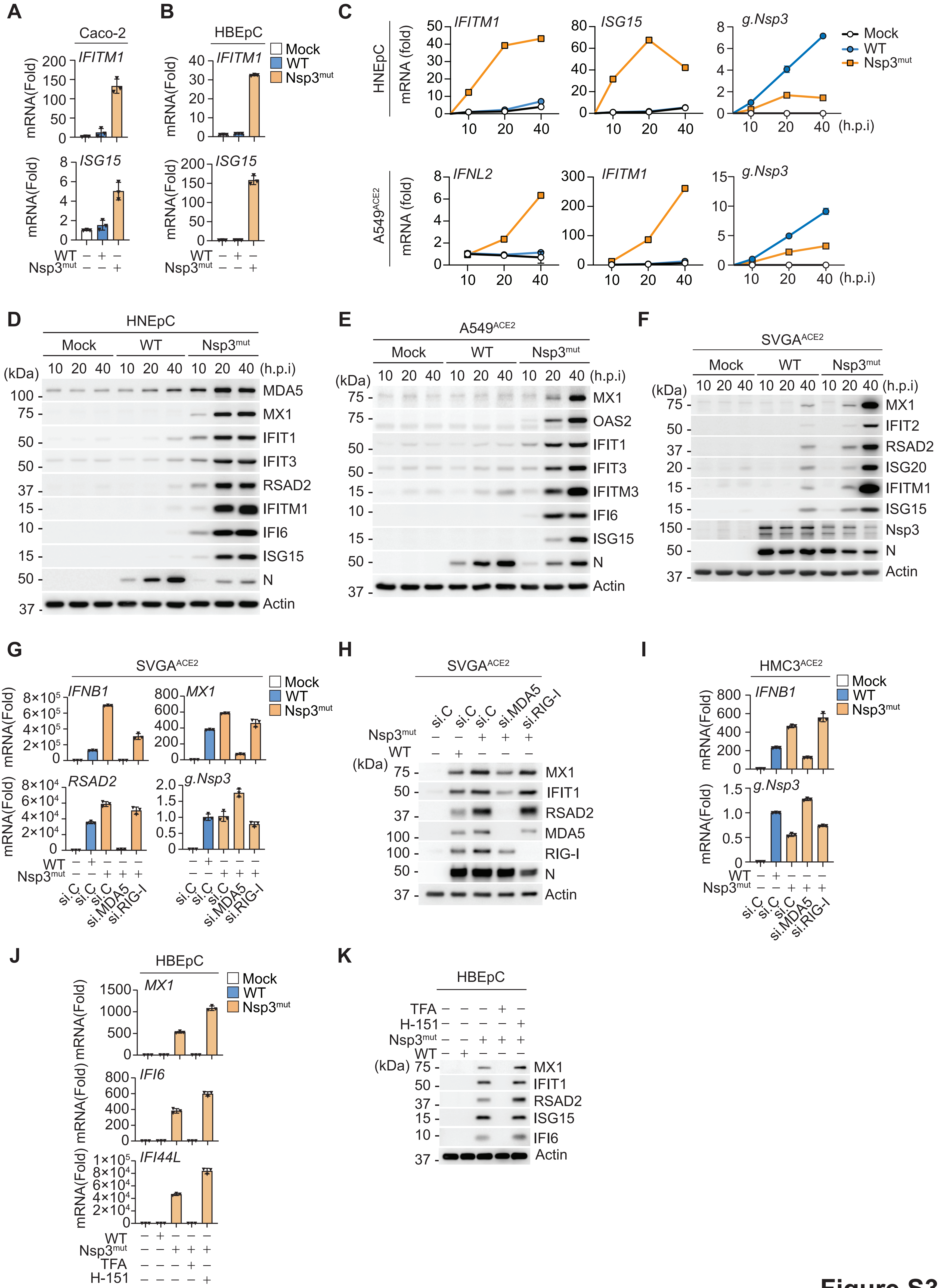

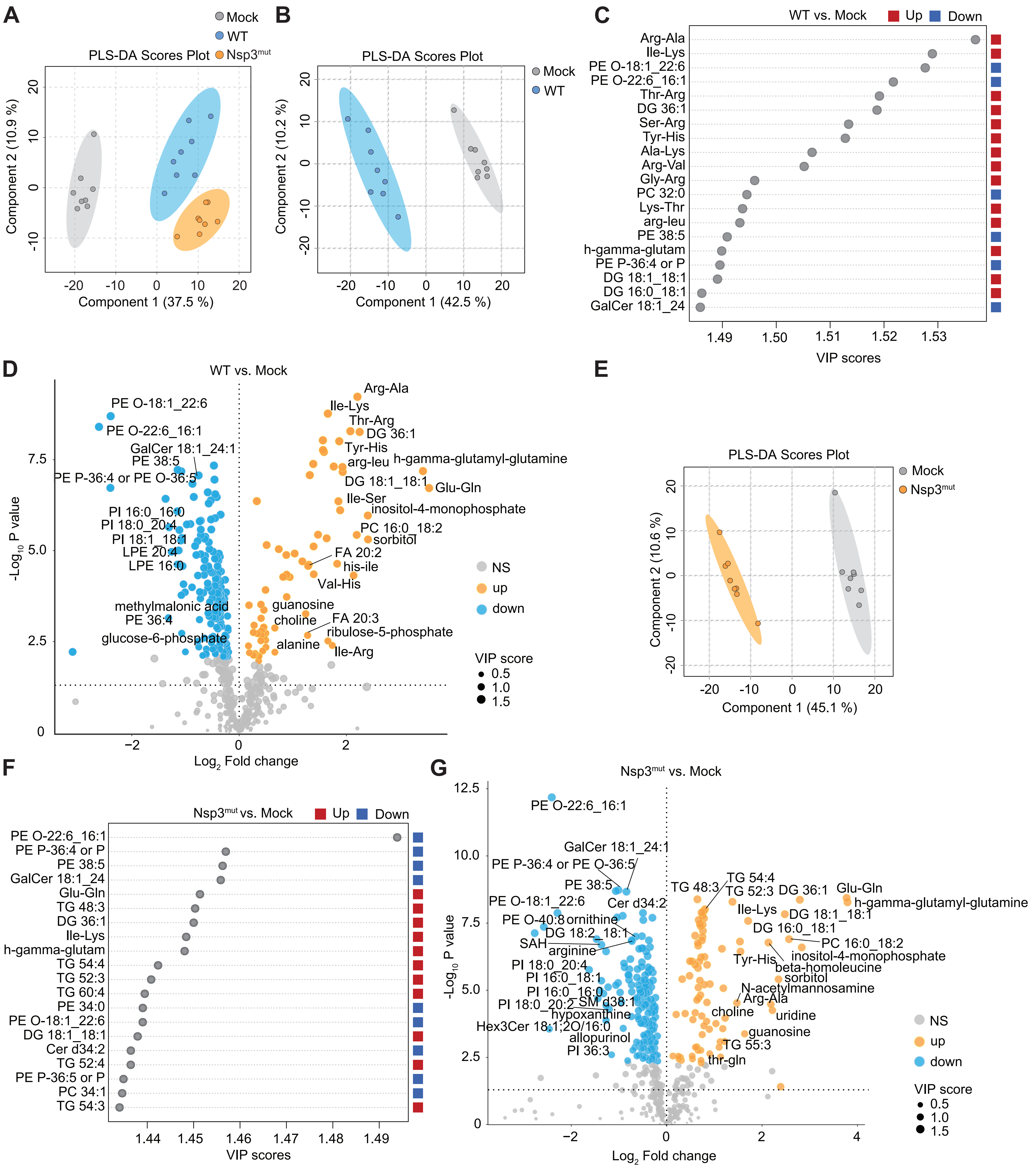

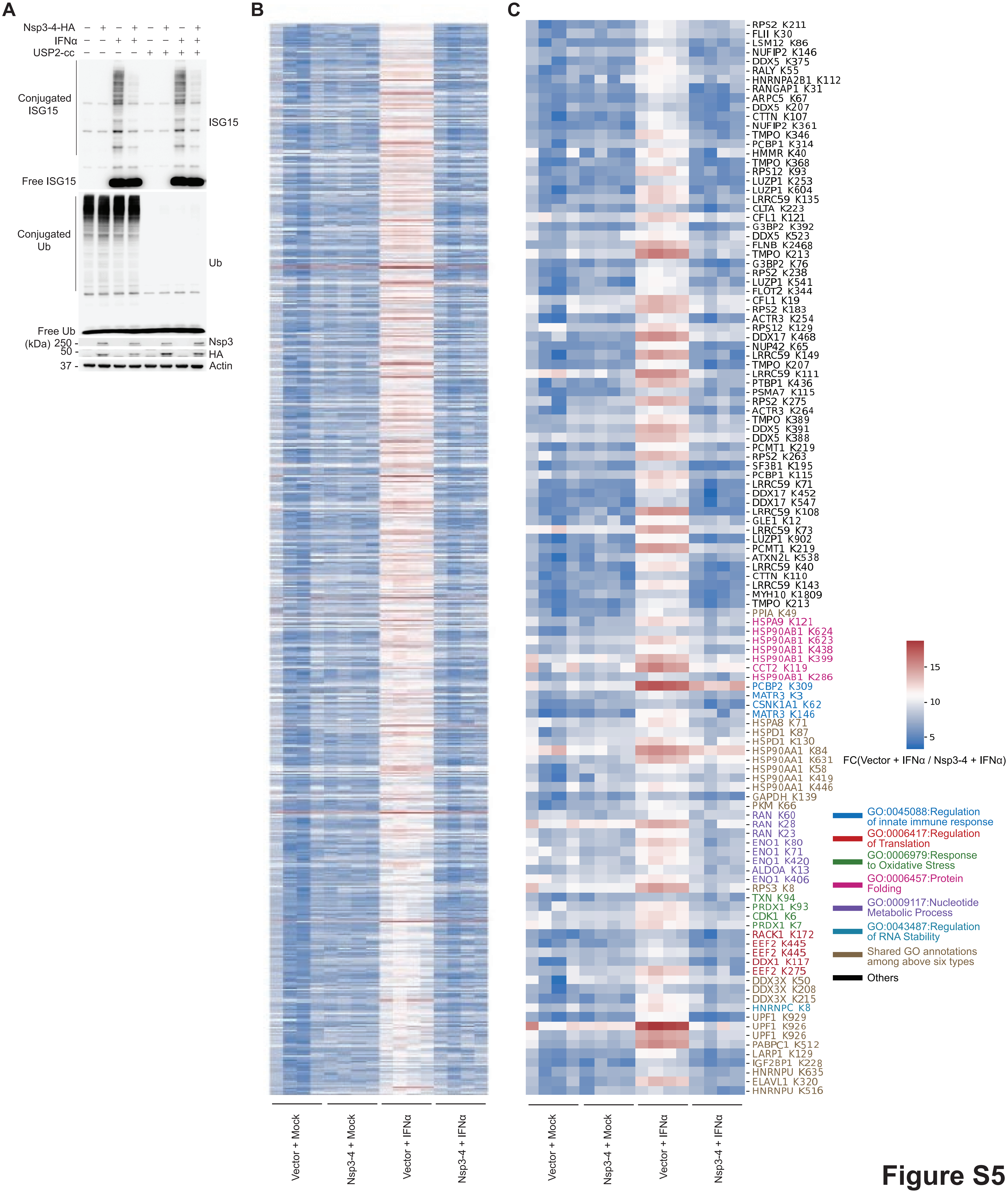

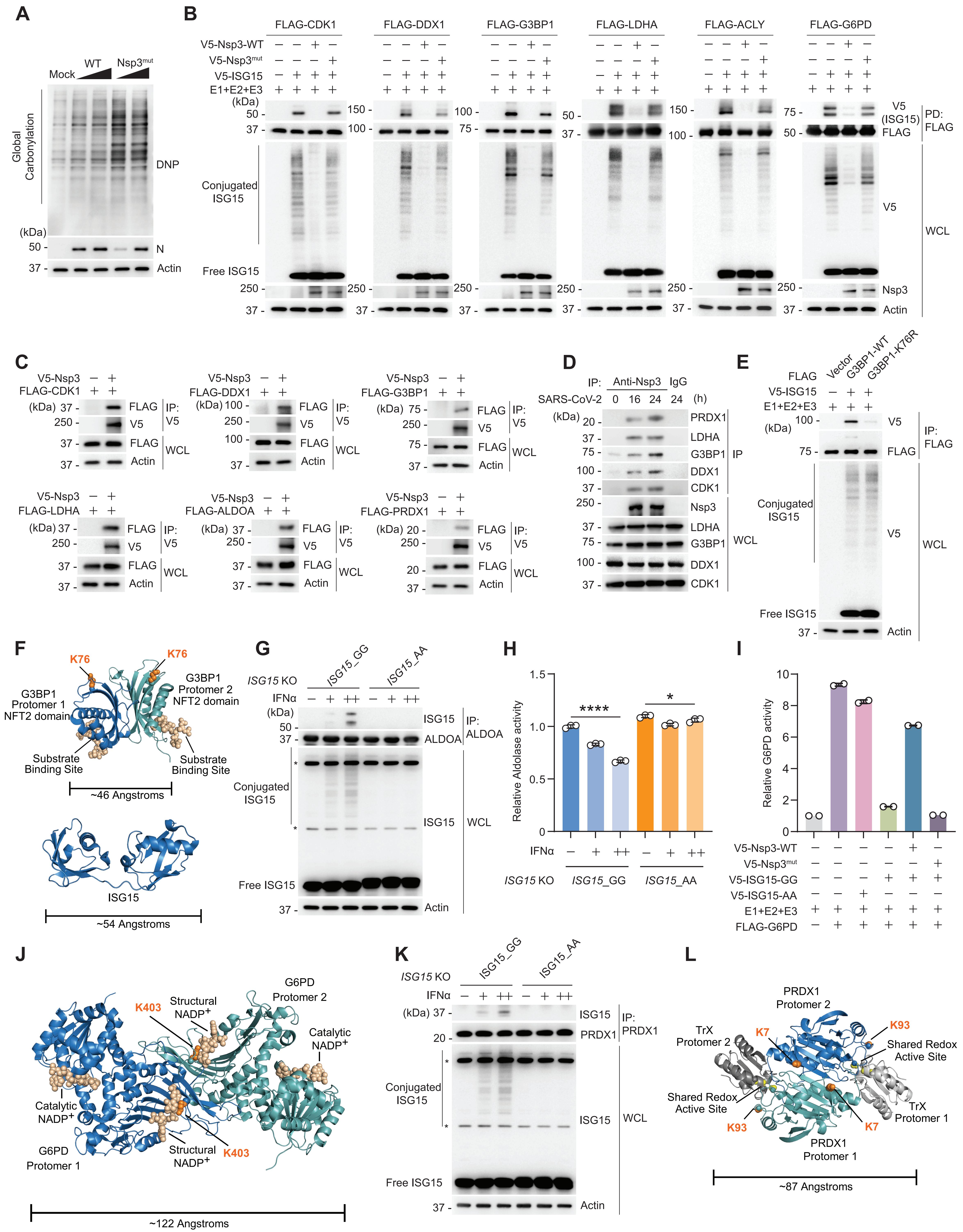
